# Antigen-driven clonal selection shapes the persistence of HIV-1 infected CD4^+^ T cells *in vivo*

**DOI:** 10.1101/2020.12.01.402651

**Authors:** Francesco R. Simonetti, Hao Zhang, Garshasb P. Soroosh, Jiayi Duan, Kyle Rhodehouse, Alison L. Hill, Subul A. Beg, Kevin McCormick, Hayley Raymond, Christopher L. Nobles, John Everett, Kyungyoon J. Kwon, Jennifer A. White, Jun Lai, Joseph B. Margolick, Rebecca Hoh, Steven G. Deeks, Frederic D. Bushman, Janet D. Siliciano, Robert F. Siliciano

**Affiliations:** Department of Medicine, Johns Hopkins University School of Medicine, Baltimore, MD; Department of Molecular Microbiology and Immunology, Johns Hopkins Bloomberg School of Public Health, Baltimore, MD; Department for Organismic and Evolutionary Biology, Harvard University, Cambridge, MA; Department of Microbiology, University of Pennsylvania Perelman School of Medicine, Philadelphia, PA; Division of HIV, Infectious Diseases, and Global Medicine, University of California, San Francisco, CA; Howard Hughes Medical Institute, Baltimore, MD

## Abstract

Clonal expansion of infected CD4^+^ T cells is a major mechanism of HIV-1 persistence and a barrier to cure. Potential causes are homeostatic proliferation, effects of HIV-1 integration, and interaction with antigens. Here we show that it is possible to link antigen responsiveness, full proviral sequence, integration site, and T cell receptor β-chain (TCRβ) sequence to examine the role of recurrent antigenic exposure in maintaining the HIV-1 reservoir. We isolated Cytomegalovirus (CMV)- and Gag-responding CD4^+^ T cells from 10 treated individuals. Proviral populations in CMV-responding cells were dominated by large clones, including clones harboring replication-competent proviruses. TCRβ repertoires showed high clonality driven by converging adaptive responses. Although some proviruses were in genes linked to HIV-1 persistence (*BACH2*, *STAT5B, MKL1*), proliferation of infected cells under antigenic stimulation occurred regardless of the site of integration. Paired TCRβ-integration site analysis showed that infection could occur early or late in the course of a clone’s response to antigen and could generate infected cell populations too large to be explained solely by homeostatic proliferation. Together these findings implicate antigen-driven clonal selection as a major factor in HIV-1 persistence, a finding that will be a difficult challenge to eradication efforts.

## Introduction

Latently infected CD4^+^ T cells represent the main barrier to HIV-1 cure (1-3). Decay of the latent reservoir, defined as the pool of cells containing inducible and replication-competent proviruses, is slow (4, 5), necessitating life-long antiretroviral therapy (ART). Proviral sequencing and integration site analysis have shown that proliferation of infected cells is a major mechanism of HIV-1 persistence (6-11). Proliferation begins early after transmission (12) for cells carrying both defective and replication-competent proviruses (13-18).

Despite evidence for *persistence by division*, factors driving proliferation of infected cells remain unclear. Homeostatic proliferation has been linked to reservoir maintenance and size (19). Stimulation with IL-7, a key mediator of T cell homeostasis and survival (20), allows CD4^+^ T cell activation and proliferation without latency reversal (21, 22). In addition, retroviral integration in certain genes may affect cell fate by insertional mutagenesis (23). Proviruses integrated in *STAT5B* and *BACH2* are frequently detected in individuals on ART (9, 24, 25) and are clustered within specific introns in the same orientation relative to host gene transcription, suggesting post-integration positive selection. However, selection of specific integration sites has been observed only for a few of the large number of genes in which integration has been observed.

The majority of HIV-1-infected cells are found in central and effector memory CD4^+^ T cell subsets (19, 26, 27), which rely on highly regulated programs of proliferation and differentiation driven by antigen stimulation (28, 29). Longitudinal analysis of the HIV-1 reservoir during ART shows fluctuating dynamics of infected cells resembling the typical expansion-contraction phases of adaptive immune responses (30). Proviral DNA has been detected in CD4^+^ T cells specific for Herpesviruses (31, 32), *Candida albicans (33)*, influenza (34), tetanus (34, 35) and HIV-1 itself (36). However, it is unclear whether enrichment of HIV-1-DNA in certain antigen-specific cells is due to higher susceptibility to HIV-1 infection (36) or higher proliferation of infected cells driven by recurrent antigenic exposure (37). Recently, Mendoza *et al*. showed that CD4^+^ T cell clones containing either defective or intact proviruses are present among CD4^+^ T cells responsive to chronic viral antigens (38). To provide definitive evidence for antigen-driven clonal expansion *in vivo* and assess its contribution to HIV-1 persistence relative to integration site driven proliferation, we studied the adaptive CD4^+^ T cell responses to CMV and HIV-1 Gag in individuals on ART.

## Results

### Accessing HIV-1-infected antigen-responsive CD4^+^ T cells

To study antigen-responding CD4^+^ T cells infected with HIV-1, obtained peripheral blood mononuclear cells (PBMCs) from 10 HIV-1-infected adults on suppressive ART for a median of 8 years (Table S1). Cells were depleted of CD8^+^ T cells and stimulated with CMV or HIV-1 Gag antigens for 18 hours. Because the CMV peptidome is large (213 ORFs) and CMV-specific T cell responses are broad and heterogeneous (39, 40), we used lysates of CMV-infected cells rather than immunodominant proteins (e.g. pp65). To study HIV-1-specific cells, we used overlapping Gag peptides since responses to class II-restricted Gag epitopes are well characterized (41). Responding cells were sorted based on upregulation of both CD40L and CD69 (Figure 1A). Median frequencies of CMV- and Gag-responding cells among all CD4^+^ T cells were 2.5 and 1.8%, respectively (Figure 1B), comparable to previous studies (40, 42). For each antigen, we also sorted CD40L and CD69 double negative cells with high expression of CD45RO to obtain memory populations depleted of responding cells (Figure 1A). PBMCs from HIV^−^CMV^−^ donors showed no increase in CD40L^+^CD69^+^ cells compared to conditions without antigens (Figure 1B and Supplementary Figure S1C). We also sorted CD40L^+^CD69^+^ responding to anti-CD3/anit-CD28 beads to obtain a population representing the CD4^+^ T cell repertoire (Figure 1A). As expected, a higher proportion of cells became activated in response to anti-CD3/CD28 beads (median 30%, p<0.0001) and a lower fraction of these were memory cells (Figure S1B).

**Figure 1.**
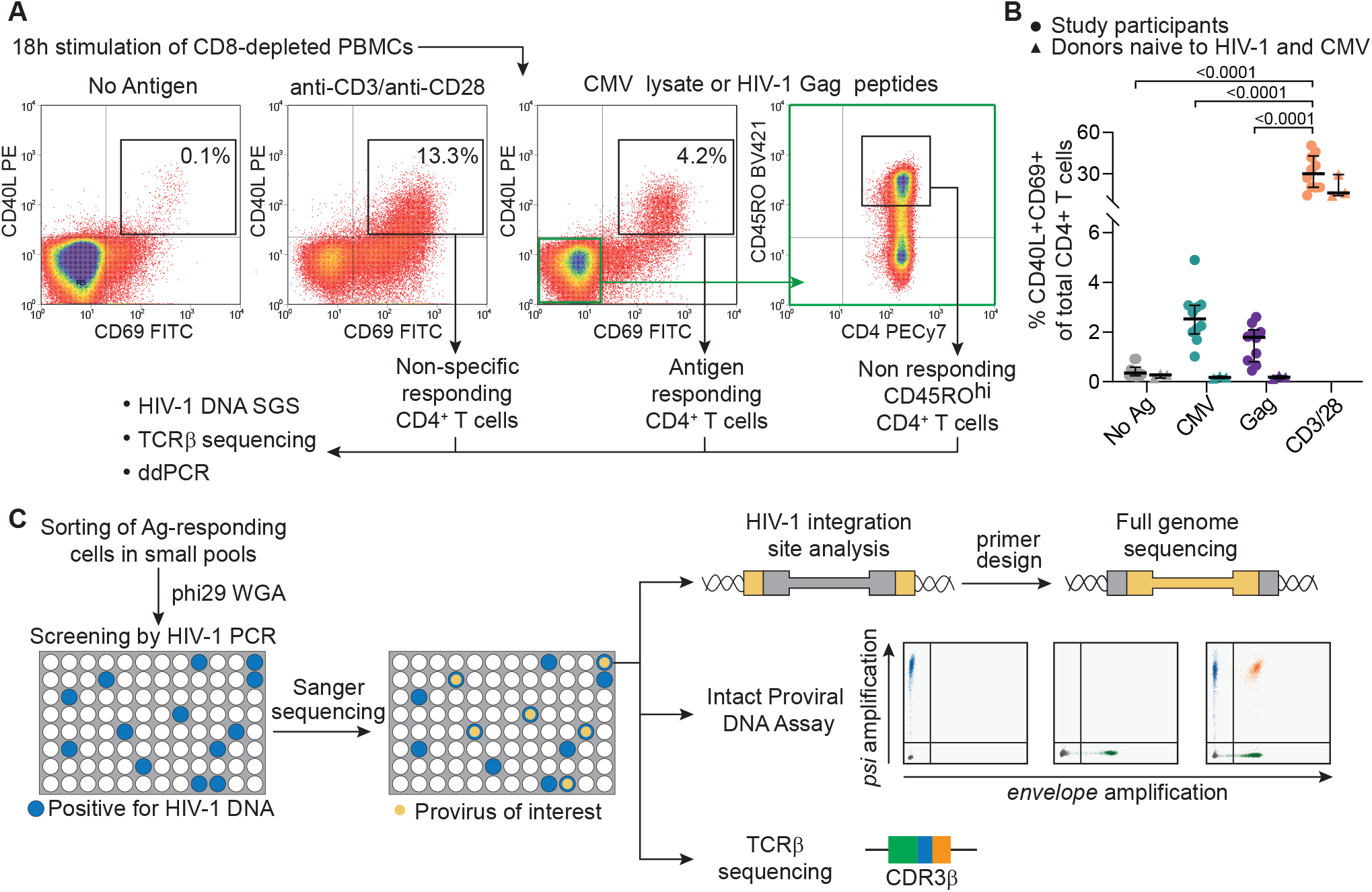
Experimental approach to study HIV-1-infected antigen-responding CD4^+^ T cells. (**A**) Experimental design and gating logic to isolate cells responding to stimulation; CD8-depleted PBMCs are stimulated with no antigen, anti-CD3/CD28-conjugated beads, CMV lysates or HIV-1 Gag peptides. CD4^+^ T cells upregulating both activation markers CD40L and CD69 were sorted. For CMV and Gag stimulations, non-responding cells with high CD40RO expression were also isolated (highlighted in green). (**B**) Frequencies of CD4^+^ T cells responding to the indicated stimulation; mean values of all timepoints are showed for each of 10 participants; horizontal bars show median and interquartile values. Statistical significance was determined by one-way ANOVA test. (**C**) Experimental design to characterize clones of HIV-1-infected antigen-responding cells. Samples from follow up time points were processed as in (A); responding cells are sorted in small pools and subjected to whole genome amplification. Pools containing infected cells are detected by u5-gag or env PCR. Proviruses matching potential clones previously identified by single genome sequencing, are detected by Sanger sequencing. Whole genome amplified DNA is then used for integration site analysis, full proviral genome sequencing, the intact proviral DNA assay and TCRβ sequencing.

Initially, DNA from sorted cells was used for single genome sequencing of HIV-1 proviruses to study the clonality of infected cells, while TCRβ sequencing was used to assess the clonality of all sorted cells (see Methods). Subsequently, responding cells from the same donors (Figures S1) were sorted in small pools representing limiting dilution with respect to infected cells (Figure 1C). Whole genome amplification (WGA) with phi29 polymerase was then used to generate thousands of copies of cell genomes (43, 44) (Figure S3). PCR on WGA-DNA identified HIV-1-positive pools, and then sequencing identified proviruses of interest for determination of integration site and full proviral sequence (Figure 1C).

### Identical proviral sequences are common among Ag-specific cells

Figure 2A shows a phylogenetic tree of 186 HIV-1 sequences from independent limiting dilution PCRs from a representative participant (P2). A higher frequency of identical sequences (appearing as “rakes” on the tree) was present in both antigen-responding (0.65 and 0.67 for CMV and Gag, respectively) and unrelated memory cells (0.68) compared to cells responding anti-CD3/CD28 stimulation (0.21), likely reflecting the presence of naïve cells in the latter condition. We identified at least one set of identical sequences in Ag-responding cells from all participants (10/10 for CMV and 8/8 for Gag, Figure 2B, Figure S4, and Table S2). Integration site analysis proved that most identical proviral sequences were true clones of infected cells (see below). In aggregated single genome sequences from all participants (n=1787), a higher fraction of the proviral sequences were identical among CMV-responding cells than among Gag-responding cells, anti-CD3/CD28 responding cells, or memory cells that did not respond to CMV or Gag (Figure 2B). These results link responsiveness to a chronic viral antigen to *in vivo* proliferation of HIV-1-infected cells. Figure 2C shows clonal proviruses with defined integration sites (see below) dominating HIV-1-infected, CMV-responding cells in four participants (P1, P3, P5, P8); these sequences were identified in multiple samples collected up to 10 months apart, demonstrating stable persistence.

**Figure 2.**
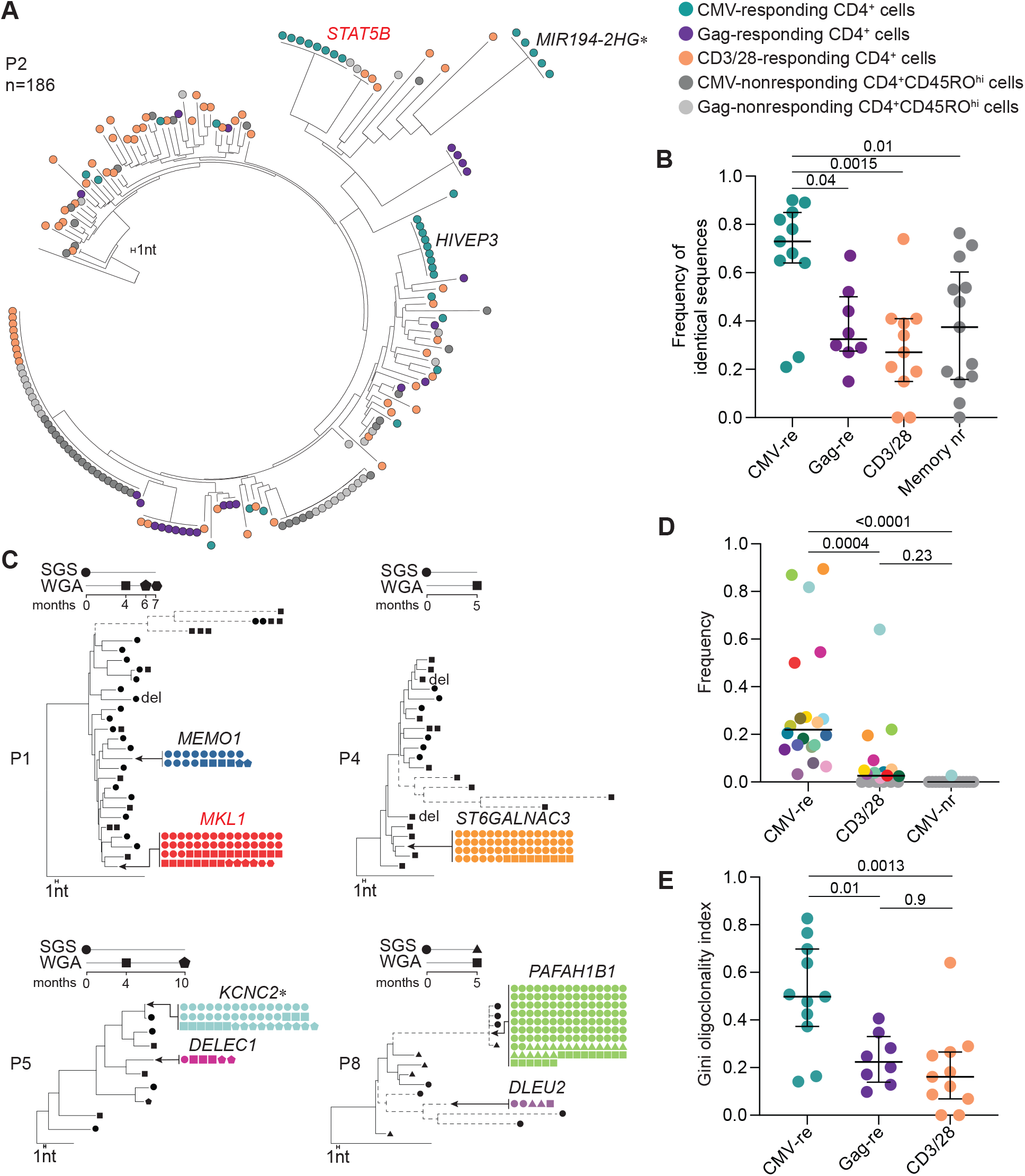
HIV-1-infected CMV-responding cells are enriched in proviral populations generated by clonal expansion. (**A**) Representative neighbor-joining (NJ) tree of 186 independent HIV-1 u5-gag DNA single genome sequences (SGS) from participant P2, rooted to HIV-1 subtype C consensus. Sequences from different sorted populations are color-coded (legend). A branch distance of 1 nucleotide is shown on the tree scale. (**B**) Frequencies of identical proviral sequences within sorted populations for all 10 participants. (**C**) NJ trees of HIV-1 sequences recovered from CMV-responding CD4^+^ T cells from 4 participants. Identical sequences are collapsed onto the same branch, and trees are rooted to the HIV-1 subtype B consensus sequence. Dashed branches indicate hypermutated proviruses. Symbols indicate method and timepoint used to generate sequences. Large CMV-specific clones are colored as in (D) and the gene containing or closest to the integration site is indicated (see Table S3 for detailed integration site data). (**D**) Dot plot showing increased frequencies of probable clones identified in CMV-responding cells compared to cells responding to CD3/CD28 stimulation or CMV-nonresponding memory cells. Only clones confirmed by integration site or potential clones comprised of at least 4 sequences were included. Probable clones are color-coded across stimulation conditions and as in (C) and Figure S4. (**E**) Dot plot showing higher clonality of proviral populations from CMV-responding cells measured with Gini coefficient. Horizontal bars show median and interquartile range. Horizontal bars show median and interquartile range.

Considering all sets of 4 or more identical sequences as potential clones, we calculated the frequencies of cells from each potential clone among all HIV-1-infected, CMV-responding cells from each donor. Potential clones had a median frequency of 0.22 (range 0.03-0.89). In the control population of CMV-nonresponding CD45RO^hi^ cells, the same sequences were either absent or present at a much lower frequency (Figure 2D). In some cases, the relative abundance of these clonal variants was not only dominant among HIV-1-infected, CMV-responding cells, but also among HIV-1-infected cells responding to anti-CD3/CD28 (Figure S4), suggesting that in some cases, CMV-specific clones represent the most expanded clones of HIV-1-infected cells. For example, in participant P8, sequences from the provirus integrated in the *PAFAH1B1* gene represented 87% of all HIV-1 sequences from CMV-responding cells and 22% of those from cells responding to anti-CD3/CD28 (Figure 2C and S4). To compare the clonality of HIV-1-infected cells across conditions, we used the Gini coefficient, a measure of distribution previously used to estimate oligoclonality in HTLV-1 infection (45). Proviral populations from CMV-responding cells showed significantly higher oligoclonality than those from Gag-or anti-CD3/28-responding cells (Figure 2E). As expected, non-responding memory cells also contained groups of identical sequences (Figures 2A and S4), specific for other unknown antigens and in some cases present at high frequencies.

These results show that clonally expanded proviral populations are present within CMV- and Gag-responding CD4^+^ T cells; the significantly higher clonality of infected cells responding to CMV suggests that the chronic immune responses to CMV antigens, characterized by memory inflation (31), contribute to proliferation and maintenance of the HIV-1 reservoir in many infected individuals.

### Integration site analysis of HIV-1-infected, antigen-responding clones

To confirm that the identical proviral sequences result from clonal expansion, we recovered integration sites using linker-mediated PCR (LM-PCR) on whole genome amplified DNA from pools of antigen-responding cells that contained identical proviruses (Figure 1C). We retrieved integration sites of 22 expanded clones, 16 from CMV-responding cells and 6 from Gag-responding cells (Figure 3A and Table S3). Most were within introns (19/22), consistent with previous studies (46, 47). There was no bias in orientation relative to the host gene (9 in same and 13 in opposite orientation). Ten integration sites were in cancer-related genes; among these, we found one provirus in *MKL1*, four in *BACH2* and two in *STAT5B*. HIV-1 integration has been previously described in these 3 genes (Table S3). Integration in *BACH2* and *STAT5B* has been linked to HIV-1 persistence due to gene activation by promoter insertion (23). The proviruses identified here shared the same features of those previously found in individuals on ART (Figure 3B). Strikingly, all four proviruses in *BACH2* were in the same orientation relative to host gene transcription and upstream of the *BACH2* translation start site, similar to 55 *BACH2* integrants identified previously. Moreover, despite defects including deletions and/or inversions (Figure 3A,), these proviruses retained the 5’ LTR and splicing donor sequences required to generate LTR-driven chimeric transcripts (48, 49). Similarly, the two proviruses in *STAT5B* were in the same orientation as *STAT5B* in intron 1, upstream of the translation start site, as with 42 of 57 proviruses previously described. These unique features, likely the result of post-integration positive selection, suggest that some HIV-1-infected clones not only proliferate in response to antigen, but also gain a survival advantage from effects of HIV-1 integration. However, 15/22 proviruses from Ag-responding expanded clones showed integration sites in loci not associated with proliferation of HIV-1 infected cells (Table S3). Moreover, some of these were near or within genes with only trace to low mRNA levels in lymphocytes, as their expression is restricted to other tissues (see Methods). CD4^+^ T cell clones carrying these proviruses have likely undergone extensive proliferation with negligible contribution of HIV-1 integration-related effects.

**Figure 3.**
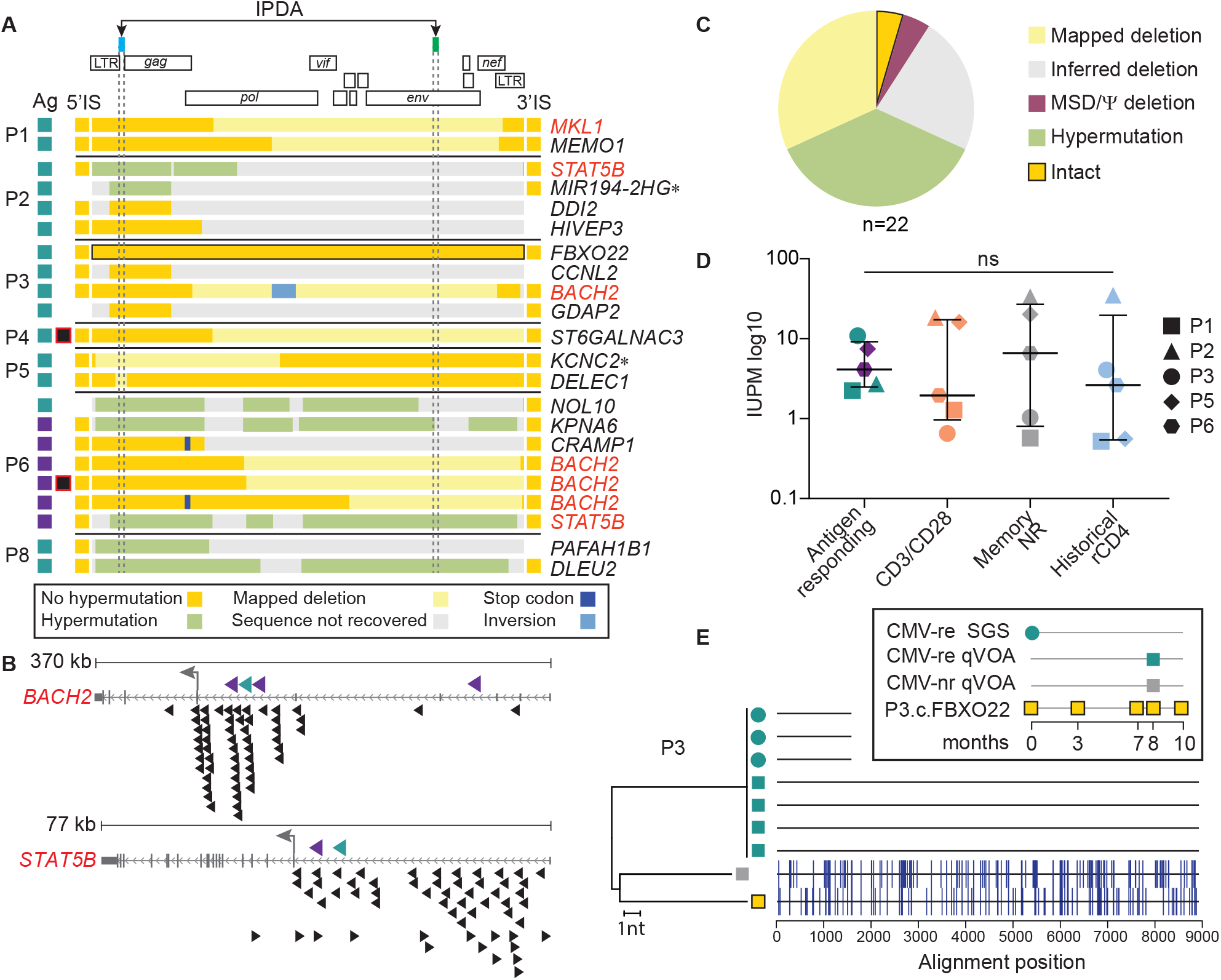
Characterization of defective and infectious proviruses from antigen-responding CD4^+^ T cell clones. (**A**) Genome sequences and integration sites recovered from proviruses in Ag-responding clones, from each participant. Each horizontal bar represents one provirus found in CMV- or Gag-responding cells (indicated by teal or purple boxes, respectively). Sequence features are color-coded according to the boxed legend. The intact provirus from participant P3 is highlighted in black. Captured host-proviral junctions are depicted as squares flanking the horizontal bars and gene symbols listed on the right show the gene containing or closest to (*) the integration site (IS); genes previously linked to the persistence of HIV-1 infected cells are highlighted in red. Proviruses with asymmetrical aberrant integration are marked with black and red squares. (**B**) Integration sites in BACH2 and STAT5B found in CMV- and Gag-responding clones (teal and purple, respectively) compared to those previously reported in individuals on ART (black). Arrow direction represents proviral orientation relative to host gene transcription. Small grey arrows show host gene transcriptional orientation, large gray arrows show translation start site. (**C**) Summary of proviral sequences in (A). (**D**) Frequency of infected cells carrying inducible replication-competent proviruses from 5 participants. Horizontal bars show median and interquartile values. Statistical significance was determined using a one-way ANOVA. (**E**) Neighbor-joining tree including u5-*gag* sequences from p24-positive qVOA wells, genomic DNA SGS and provirus P3.c.FBXO22, sampled according to the legend in the insert. The highlighter plot shows mismatches from the top sequences in the tree.

### Antigen-responding clones carry both defective and infectious proviruses

To investigate whether clones of HIV-1-infected, antigen-responding cells carried intact or defective proviruses, we recovered the partial or full-length sequences of the proviruses for which we identified integration sites (Figures 3A and C). As expected, most genomes (13/22) were defective due to mapped or inferred deletions or G-to-A hypermutation (see Supplementary Data). However, we did detect 1 intact genome, Despite the limited sampling, this proportion of intact and defective proviruses was consistent with previous analysis of the proviral landscape in CD4+ T cells (50).

We also carried out quantitative viral outgrowth assays (51) (qVOAs) (Figures 3D and E and S7) on the sorted cell populations (and observed outgrowth from all populations from all participants. Infected cell frequencies in infectious units per million cells (IUPM) were not significantly different for CMV- or Gag-responding cells than for cells responding to CD3/CD28 stimulation or for non-responding memory cells; in addition, there was no difference relative to historical qVOAs using resting CD4^+^ T cells from the same individuals (Figure 3D), suggesting that, at least in these participants, the HIV-1 reservoir was not enriched in cells responding to the antigens tested.

To determine whether the induced infectious proviruses were from clonally expanded cells, we sequenced the HIV-1 RNA from the supernatants of p24 positive wells (Figure 3E and S7). In participant P3, we detected viral outgrowth in four independent wells seeded with CMV-responding cells (corresponding to 10.8 IUPM). These replication-competent viruses had identical near-full length genome sequences, which were genetically distinct from one outgrowth virus recovered from CMV-nonresponding cells (1.03 IUPM). In addition, these four identical isolates matched 3 HIV-1 DNA single genome sequences found in CMV-responding cells 8 months previously. Interestingly, outgrowth of the intact provirus integrated in the *FXBO22* gene, which was the most abundant sequence among HIV-1-infected CMV-responding cells from participant P3 (frequency of 0.2, Figure S4) and persisted across multiple time points, was not detected (Figure 3E), suggesting lack of inducibility or the presence of missense mutations that would affect replicative fitness.

These results provide strong evidence that CD4^+^ T cell clones carrying an inducible replication-competent provirus can be selected over time in response to a CMV. However, the size of infectious proviral clones within antigen-specific populations varied, as in the study from Mendoza *et al*. (38). In participant P2, cells responding to anti-CD3/CD28 and CMV-nonresponding cells had a significantly higher frequency of cells carrying inducible, replication-competent proviruses (18.3 and 33.5 IUPM, respectively, versus 2.7 IUPM in CMV-responding cells). This was likely due to a single large clone specific for an antigen other than CMV (Figure S4). Proviral sequences identical to this outgrowth virus were found in PBMCs from 8 months previously, when they represented the most abundant viral variant in both CD3/CD28 and CMV- or Gag-nonresponding memory cells (frequency of 0.15, 0.49 and 0.32, respectively, Figures 2A and S4). Thus, other factors, including antigens not explored here, can lead to massive expansion of clones carrying infectious proviruses.

### TCRβ repertoire mirrors the clonality of proviral populations

VDJ recombination generates a T cell receptor (TCR) repertoire with a theoretical diversity exceeding the total body number of T cells (52) (up to 10^15^). Therefore, we used VDJ rearrangements as T cell barcodes to compare clonality in CMV- and Gag-responding cells with that in non-responding memory T cells and cells activated by anti-CD3/CD28. Despite comparable sequencing depth (Figure S8), species richness for TCRβ sequences in CMV- and Gag-responding cells was significantly lower than for CD3/CD28 responding cells (Figure 4A), and the oligoclonality indices for CMV-responding cells were not only higher than those of CD3/CD28 stimulated cells, but also than those of Gag-responding cells (Figure 4B). Thus, patterns of clonality for the whole responding CD4^+^ T cell population mirrored those of the HIV-1-infected cells within that population. Oligoclonality of TCRβ sequences correlated with oligoclonality of proviruses within the relevant populations (Spearman coefficient r=0.52, p=0.01), and samples from CMV-responding cells clustered away from anti-CD3/28 and Gag-responding cells (Figure 4C), suggesting that adaptive immune responses globally affect the expansion of HIV-1-infected clones and supporting previous studies describing inflation of memory cell populations driven by CMV (31).

**Figure 4.**
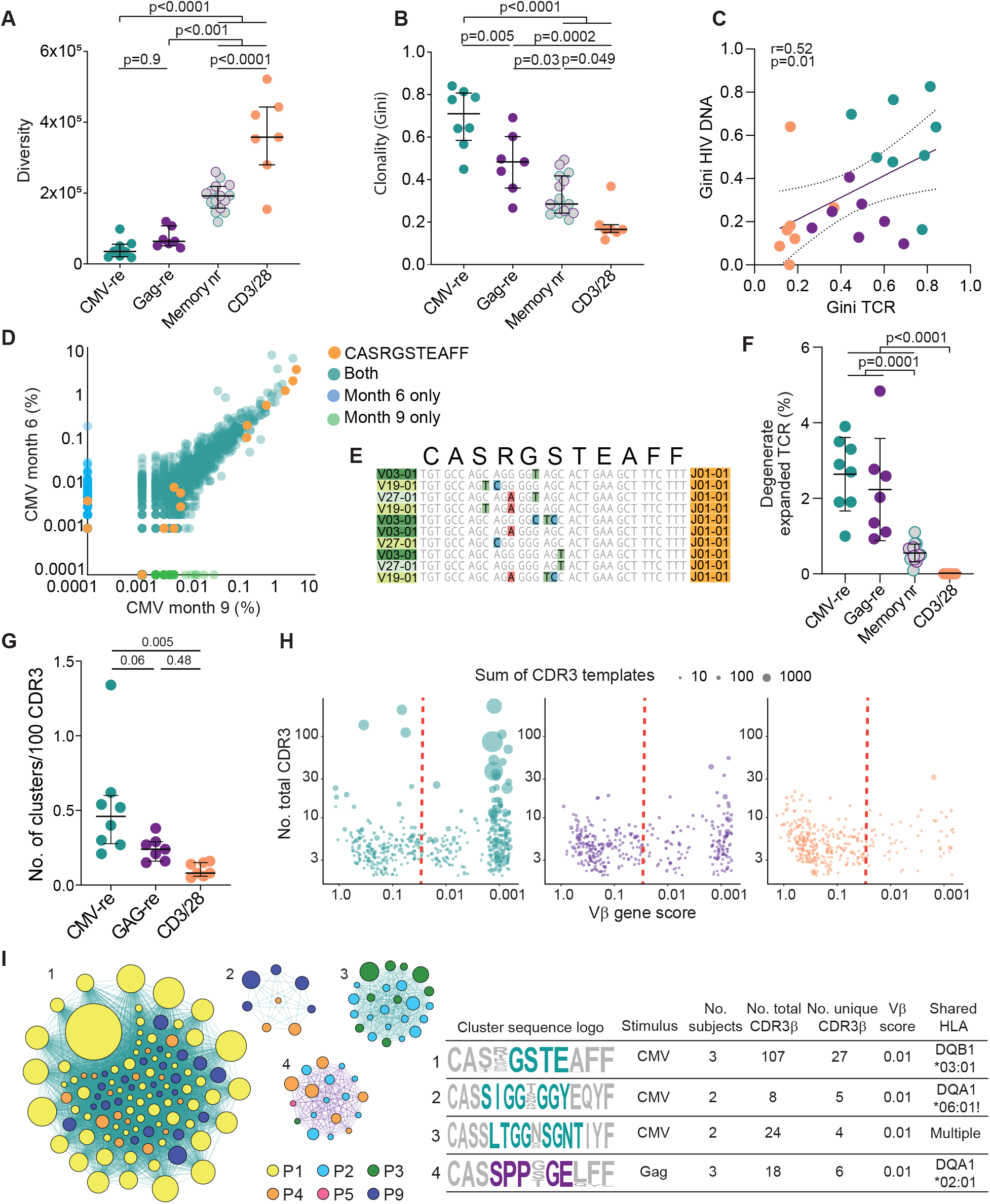
Antigen-responding cells show higher clonality and evidence of convergent selection. (**A**) TCR diversity, estimated by the Chao index (107), is lower in antigen-responding cells. Grey circles represent memory cells non responding to either CMV or Gag stimulation (teal and purple border, respectively). (**B**) Gini coefficients based on TCRs from CMV-responding cells show the highest clonality. (**C**) Correlation of Gini coefficients based on TCR and proviral populations. Spearman test and linear correlation with 95+ confidence intervals are shown. (**D**) Log10 abundance of productive TCRs from CMV-responding cells from participant P1 collected at month 6 and month 9 of the study. (**E**) VDJ sequences from the 10 most abundant clonotypes encoding CASRGSTEAFF. (**F**) Percentage of the sum abundance of degenerate and expanded CDR3β sequences. (**G**) Frequency of TCR clusters normalized by CDR3β input. Clusters were filtered based on CDR3β ≥2 and a Fisher’s exact test p<0.0001. (**H**) TCR clusters are plotted based on number of total CDR3β and Vβ gene score. Lower scores indicate more homogeneous Vβ. Circle size is scaled to the total sum of TCR templates in each group, indicating clonal expansion of cells within the cluster. (**I**) Representative TCR clusters involving more than one participant, displayed as networks and CDR3β sequence logos. Nodes represent each CDR3β sequence, with circle colors based on participant and circle size representing clonal expansion. Edge colors highlights antigen stimulation (teal for CMV and purple for Gag). CDR3β logos display amino acid representation at each position. The core motif shared by the convergent cluster is colored. The table shows cluster characteristics and shared HLA alleles (see Table S4 for additional details). Exclamation marks indicate enrichment of HLA allele in participants contributing to the cluster. Horizontal bars show median and interquartile values. Statistical significance was determined using a one-way ANOVA.

To estimate the distribution and the relative sizes of CMV- and Gag-responding T cell clones within the entire CD4^+^ T cell repertoire, we identified TCRβ sequences with a significantly higher abundance in CMV- and Gag-responding cells than in cells activated by anti-CD3/CD28 stimulation (see Methods). Across seven participants, we identified an average of 210 (SD ± 103) and 137 (SD ± 82) T cell clones enriched in CMV- and Gag-specific cells, respectively (Figure S8). Of note, a significantly smaller number of clones were enriched upon either stimulation (average 16, SD ±10, p=0.0004), likely reflecting cross-reactivity or non-specific activation. Interestingly, although these CMV- and Gag-reactive T cell clones represented only a small percentage of all CD4^+^ T cells (average of 2.5 and 1%, respectively, Supplementary Figure S8C and D), they were among the top expanded T cell clones in the anti-CD3/CD28 responding population. CMV-responding T cell clones were particularly dominant (Figure S8), as was the case in the analysis of clonal proviral populations among HIV-1-infected cells (Figures 2D and S4).

To prove the striking clonality of CMV-responding CD4^+^ T cells resulted from antigen-dependent selection, and not homeostatic proliferation, we examined the most expanded clonotypes (defined here as cell clones sharing an identical VDJ β sequence) and found many shared the same amino acid sequence in the CDR3β region, despite different VDJ rearrangements at the nucleotide level. These so called “degenerate” TCRs are signatures of convergent immune responses, selected over time for binding to specific peptide-loaded MHC molecules (53, 54). Figures 4D and 4E show one such degenerate TCR. TCRβ sequencing on CMV-responding CD4^+^ T cells collected 3 months apart from participant P1 showed overlapping distributions of clonal frequencies (suggesting stability over time) including 19 different clonotypes with an identical CDR3β amino acidic sequence (CASRGSTEAFF, Figure 4E).

These rearrangements represented the most abundant CDR3β sequences among CMV-responding cells (7 and 10%, for the two time points). Degenerate expanded clones of this kind were significantly more abundant in Ag-responding cells (median of 2.8% for CMV and 2% for Gag, respectively) compared to non-responding and anti-CD3/28 stimulated cells (median 0.5% and 0.003%, respectively, Figures 4F and S8), supporting the hypothesis that CD4+ T cell responses against these two chronic infections undergo a process of convergent selection.

To further explore clonal selection based on TCR specificity, we performed a TCR cluster analysis of shared structural features using the GLIPH2 algorithm, which groups TCRs into clusters predicted to bind the same peptide-MHC complex (55). TCRs from CMV-responding cells had a significantly higher frequency of clusters than those from anti-CD3/28 stimulated cells (Figure 4G, median 4.6 versus 0.8 clusters every 1000 TCR input sequences, p=0.005). In addition, a higher proportion of clusters from CMV-responding cells had a larger size (included a higher number of clonotypes), included expanded clones and showed a restricted V gene use (p<0.05) (Figure 4H), supporting an overall higher degree of selection towards convergent immune responses. Finally, when we used GLIPH2 on TCRs sampled from all participants (n=8 for CMV and n=7 for Gag), we found convergent clusters including CDR3s from multiple subjects, likely representing public responses against shared immunodominant epitopes. Four exemplary clusters are described in Figure 4I and Table S4. The CDR3 sequences in these clusters showed a restricted use of V and J genes, shared significant motif residues and were often degenerate. Moreover, the individuals contributing to these clusters shared one or more class II HLA alleles. Although it is not technically feasible to extend these TCR analyses to the rare subset of CMV-responding cells that harbor latent proviruses, our results strongly support the conclusion that the clonality of CMV-responding cells is the result of antigen-driven proliferation and not a homeostatic process that occurs independent of TCR specificity.

### Coupled quantification of provirus and VDJ rearrangements

To better understand the nature of Ag-specific CD4^+^ clones carrying proviruses, we developed a novel method to identify their cognate TCRβ sequences. We sequenced the TCRβ repertoire from multiple whole genome amplified cell pools containing the same provirus and searched for the unique VDJ rearrangement that recurred across all pools carrying a provirus with a specific integration site. To confirm the assignment to TCR/provirus pairs, we leveraged the extraordinarily high diversity of the T CR repertoire and adopted combinatorial statistics previously used to confidently pair α and β chains (56) (see Methods and Supplementary Figure S9A). The patterns of co-occurrence observed for assigned pairs of proviruses and TCRβ sequences indicated that they belong to the same T cell clones and do not occur together simply by chance (*P* between 10^−3^ and 10^−13^, Supplementary Figure S9B). We identified 8 unique pairs across 6 participants, 5 from CMV-responding cells and 3 from Gag-responding cells (Figure 5). T cell clones carrying specific proviruses ranked among the most abundant CMV-responding TCR sequences (Figure 5C). As expected, the TCRs of the three Gag-specific clones paired to proviruses were less abundant, in agreement with analyses of clonality of Gag-responding cells.

**Figure 5.**
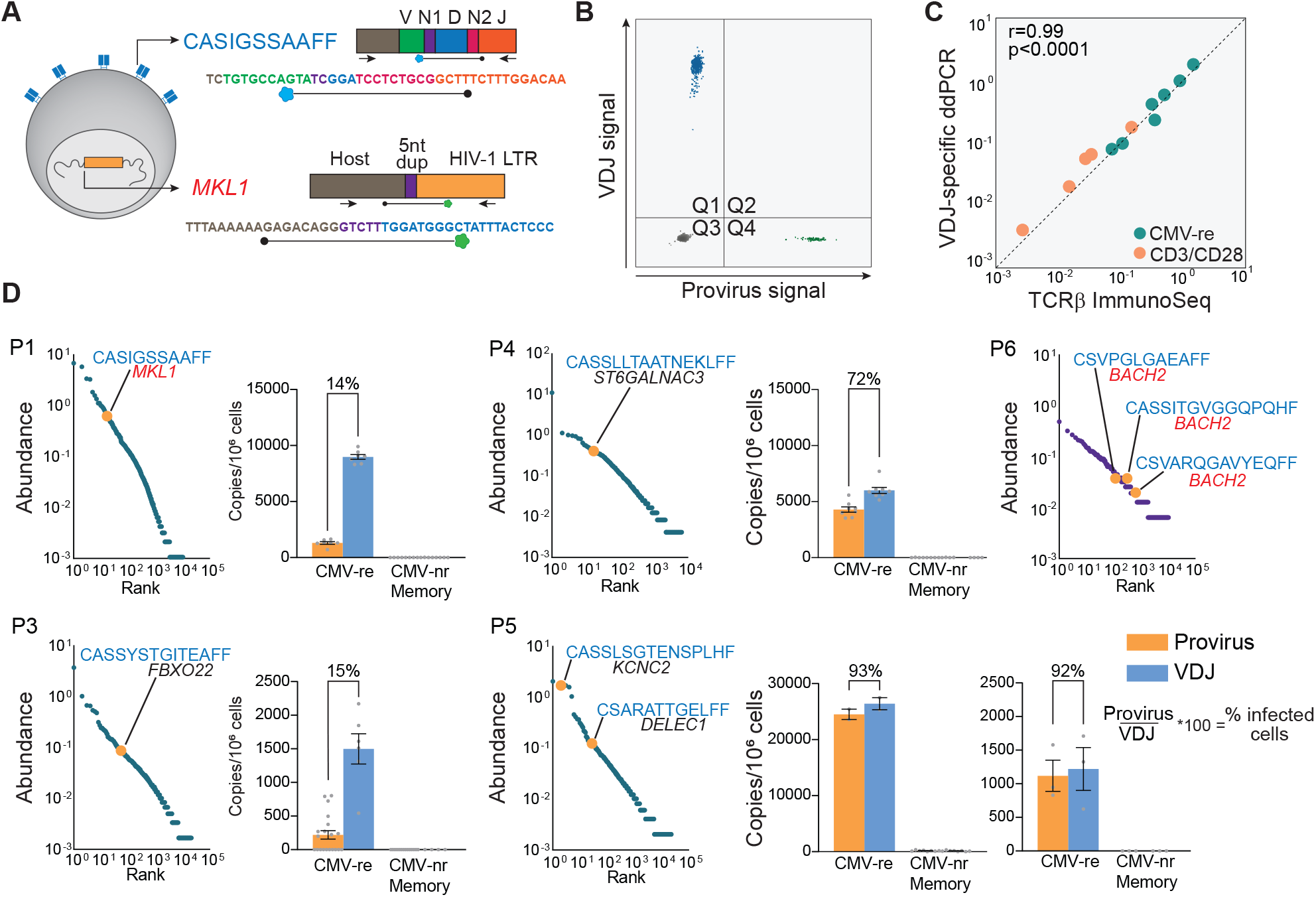
Analysis of VDJ and provirus pairs belonging to the same antigen-responding clone. (**A**) Droplet digital PCR design for duplex quantification of VDJ and proviral copies from genomic DNA of sorted cells. (**B**) Representative ddPCR 2D plot of duplex amplification of CASIGSSAAFF and cognate provirus integrated in MKL1 gene. (**C**) Quantification of clonotypes by VDJ-specific ddPCR strongly correlates with TCRβ immunosequencing; five CMV-responding clonotypes (shown in D) where quantified in sorted cells responding to CMV or CD3/CD28 stimulation; axes represent Log10 abundance. (**D**) Log10 ranked abundance plots of CMV- of Gag-responding cells showing HIV-1-infected clonotypes for which we identified both the VDJ rearrangement and the integration site, highlighted in orange. For each pair, bar graphs show the frequency of provirus (orange) and VDJ (blue) copies in CMV-responding and nonresponding memory cells. The provirus-to-VDJ ratios are used to calculate the percentage of a given clones that is HIV-1-infected.

To estimate the fraction of a given CMV-specific CD4^+^ clone that carried the associated provirus, we designed duplex digital droplet PCR (ddPCR) assays to directly count, within the same sample of sorted cells both the rearranged VDJ sequence and the host-provirus junction belonging to 5 CMV-specific clones (Figure 5A and B). These corresponding VDJ and proviral sequences were highly enriched in CMV-responding cells but absent or rare in memory cells not responding to CMV. We calculated the ratio between proviral copies (infected cells) and VDJ copies (total cells in the antigen-specific clone) (Figure 5C). We observed two patterns: in 3 clones from 3 participants, only a fraction of all the cells comprising the clone carried the proviruses (range 14-72%), suggesting that these clones were expanded before a member of the clone became infected and subsequently proliferated. Conversely, in 2 clones from participant P5, almost 100% of the cells comprising the clone carried the relevant provirus, suggesting that extensive Ag-driven clonal expansion happened after the infection event in one of the progenitor cells of that clone (Figure 6A and B).

**Figure 6.**
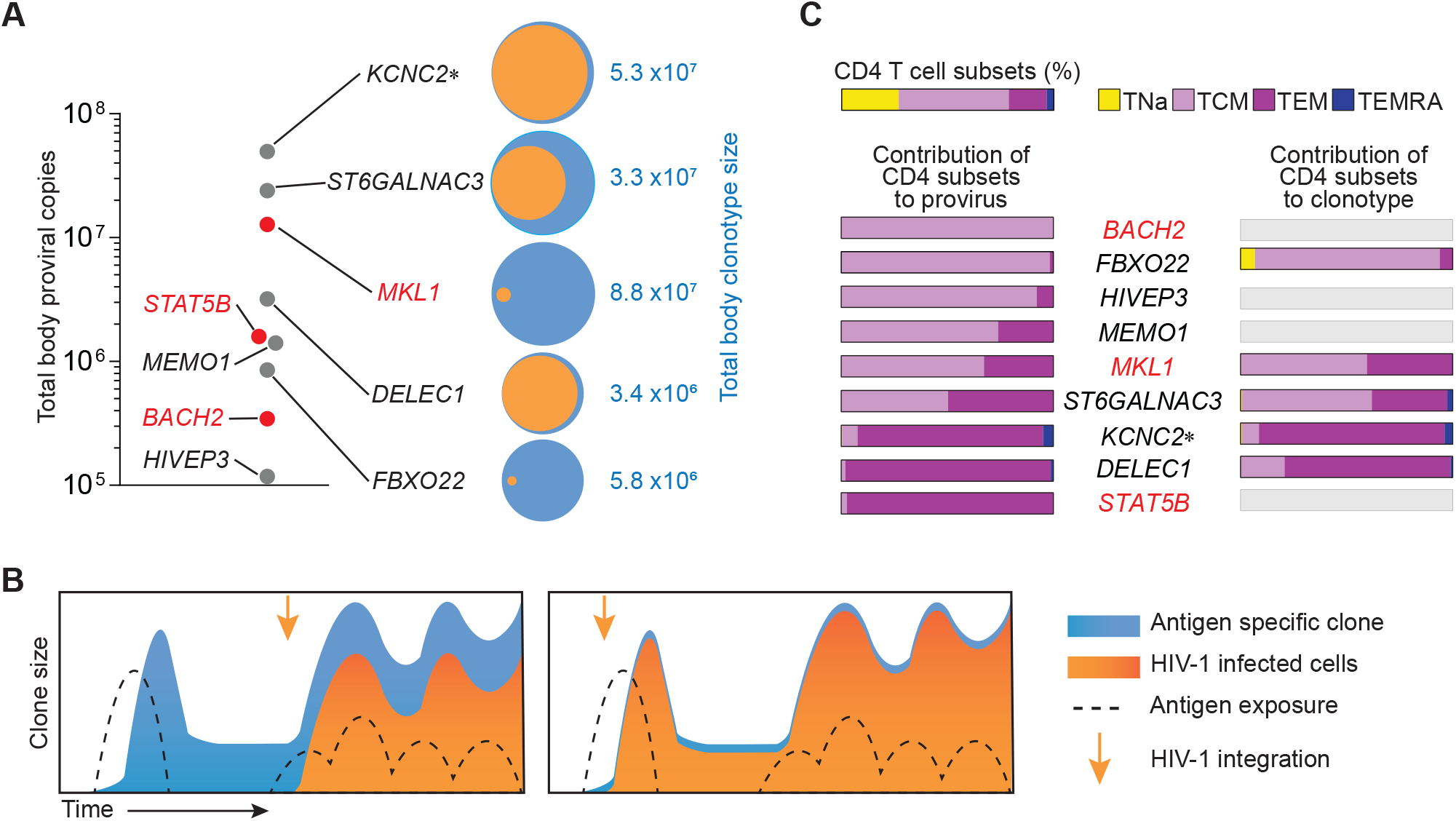
Total body clone sizes and contribution of CD4^+^ T cell subsets. (**A**) Proviral and VDJ frequencies are used to back-calculate total body clone sizes. Clones with integration sites in genes previously linked to the persistence of infected cells are highlighted in red. Venn diagrams show the fraction of a clonotype (orange) carrying its cognate provirus (blue). (**B**) Two scenarios of infection-expansion dynamics of antigen-driven clonal selection. The left panel shows infection of a clone already expanded in response to antigen, while the right panel shows selection occurring after an early infection event. (**C**) Contribution of CD4^+^ T cell memory subsets to clones based on provirus and VDJ measurements. Memory subsets are defined as shown in Supplementary Figure S10. Horizontal bars show the relative contribution of memory subsets for each clone.

To investigate the contribution of T-cell differentiation to the persistence of these Ag-specific clones, we applied the ddPCR assay to CD4^+^ T cells subsets sorted based on the expression of CCR7 and CD45RA (Supplementary Figure S10). Although we observed the expected proportions of naïve (TNa), central memory (TCM), effector memory (TEM) and effector memory CD45RA^+^ (TEMRA) cells within total CD4^+^ T cells, most proviral copies were found in TCM and TEM subsets (Figures 6C and Figure S10), in accordance with previous studies based on HIV-1 DNA (19, 26, 27), HIV-RNA (57) and on specific expanded clones carrying proviruses (58, 59). However, we observed wide variation in the relative contribution of TCM or TEM cells for different proviruses (from 0 to 100%), suggesting adaptive immune responses to individual epitopes are heterogeneous and likely influenced by other factors, like frequency of peptide-MHC stimulation, TCR affinity (60) and T-cell activation. More importantly, despite this variation, we observed a similar contribution of TCM and TEM subsets between proviral and VDJ copies of the same clone (Figure 6C), supporting the hypothesis that infected and uninfected cells within an antigen-responding T cell clone are under the same differentiation-proliferation program. This finding further supports the idea that expansion of HIV-1-infected T cell clones is mostly regulated by T cell physiology, rather than HIV-1-mediated effects.

To understand the extent of expansion of these HIV-1-infected, CMV-specific clones, we estimated their total body size. We observed striking clone sizes ranging between 10^5^ and 10^8^ cells (Figure 6A). The number of divisions needed to reach such sizes (mean 21, SD ± 2.9) is achievable, considering the short doubling time of activated T cells (61), but it does not account for cell death (during clonal contraction) and the fact that antigen-driven expansion occurs only for rare (specific) clones and not for the whole T cell population. To investigate whether clones of this size could result from homeostatic proliferation, we calculated the likelihood a clone could reach a given size by chance if the entire population of CD4^+^ T cells was maintained by a constant, balanced, process of division and death (see online methods and Table S5). For the sizes of the HIV-1-infected CMV-responsive clones described here, probabilities approached zero even for high turnover rates, supporting a scenario in which non-random events drive the proliferation of rare cells (antigen-specific clonotypes), rather than a homogeneous process of "cohort homeostasis". Thus, the major driver of infected cell proliferation and hence HIV-1 persistence, is the response to antigen and not specificity-independent homeostatic or integration-site related proliferation.

## Discussion

The requirement for lifelong treatment, due to the HIV-1 reservoir, further hinders the scalability and maintenance of HIV-1 care worldwide (62). Thus, a better understanding of the mechanisms of HIV-1 persistence is needed. Two recent studies investigated the immunological factors contributing to the proliferation of the HIV-1 reservoir: Mendoza *et al*. characterized proviral sequences from cells reactive to viral antigens (38), while Gantner *et al.* profiled TCRβ sequences from cells with inducible p24 expression (59). Here, by combining provirus, integration site and TCRβ analyses from persistent clones with known specificities, we provide new insights into the importance of antigen-driven proliferation, relative to other drivers of clonal expansion, such as integration site effects and homeostatic proliferation. Through the analysis of antigen-responding cells, we identified proviral populations resulting from clonal expansion in all participants. Some clonal proviruses were present at multiple time points for up to 25 months. In CMV-responding cells, proviral populations showed high clonality. Analyses of TCRβ repertoires confirmed the overall higher clonality in CMV-responding cells. TCRs from these cells were characterized by degenerate and often convergent CDR3β sequences, further supporting the conclusion that proliferation of infected cells is driven by adaptive immune responses affecting all cells, not just infected cells. To untangle the role of antigenic pressure relative to HIV-1 insertional mutagenesis, we determined the integration sites of infected antigen-responding clones. We identified CMV- and Gag-specific clones with proviruses in genes previously linked to the proliferation of HIV-1 infected cells (*STAT5B*, *BACH2*, *MKL1*). This is the first evidence that some HIV-1-infected cells persist with more than one selective advantage, as recently described in CAR-T cell clones (63, 64). Nevertheless, many CMV-specific clones carried proviruses in loci unlikely to affect T-cell phenotype and survival. In addition, the quantification of proviral and VDJ sequences belonging to the same clonotype indicated that in some cases only a fraction of the T cell clone is infected, suggesting antigen-driven clonal expansion before the HIV-1 integration event and subsequent proliferation of infected cells. Together, our work suggests that although proviruses can disrupt the expression of host genes and promote survival (49), the effect of HIV-1 integration is not indispensable for the expansion of antigen-specific infected cells, in accordance with a recent study on large clones carrying infectious proviruses responsible for low level viremia (65).

T cell activation reverses latency – that was how reservoir was discovered. But the extensive proliferation observed here is hard to reconcile with latency reversal, which should render cells susceptible to cytopathic effects or immune clearance. Our results further support the hypothesis is that infected cells undergo extensive proliferation without necessarily inducing viral expression (18, 66). The identification of common antigen-specific responses driving proliferation of cells in the HIV-1 reservoir could have implications for novel therapeutics strategies (37). Such antigens could reverse latency or drive differentiation of infected cells towards terminal effector subsets with shorter half-lives (67, 68). Antigens driving clonal expansion like CMV could be targeted to both reduce chronic inflammation and accelerate reservoir decay (69, 70). CD4^+^ T cells can be differentially susceptible to HIV-1 according to their specificity (71). For example, while the autocrine production of β-chemokines can protect some CMV-specific cells from HIV-1 infection (72), HIV-1-specific cells are preferentially infected during acute infection and viral recrudescence (36). Since we used a different experimental approach (activation-induced markers CD40L and CD69 instead of intracellular staining of effector molecules) and studied individuals who started ART during chronic infection and were on suppressive ART for years, we were not able to investigate whether CMV- and Gag-specific CD4+ T cells had a significantly different frequency of infected cells and confirm the observations from previous studies. However, we observed a striking heterogeneity across participants in the distribution and relative size of antigen-specific clones carrying HIV-1 genomes. Similarly, the recovery of replication-competent virus from antigen-specific cells varied for each participant and showed no reservoir enrichment, overall, in either CMV- or Gag-responding cells. Despite the small number of individuals and the limited range of antigens tested in this study, our results suggest that the specificity of HIV-1-infected cells differs extensively from person to person. In addition, our findings further support the dynamic nature of the reservoir (30). Fluctuations of clone sizes due to episodic exposure to cognate antigen can further complicate efforts to quantify the reservoir and its decay based on measurements of total or intact proviral DNA (73, 74). Future research should investigate other ubiquitous pathogens that typically co-infect people living with HIV-1 and commensal microbials that play a role in chronic antigenic stimulation. Moreover, antigen-independent mechanisms that contribute to the reservoir structure need to be explored, such as the proliferation induced by T cell depletion, that occurs upon ART introduction (75) or post-therapeutic ablation (32, 76). Similarly, steady-state homeostatic proliferation, which allows the persistence of long-term memory in the absence of antigen, likely plays a role in the maintenance of the HIV-1 reservoir.

In conclusion, our work provides a better understanding of the complex forces driving clonal expansion, showing that antigens can shape the fate of infected cells independently from the effects of HIV-1 integration. Antigen-driven clonal expansion and selection cause HIV-1-infected clones to persist indefinitely, proliferate, and reach sizes that make eradication daunting, except with strategies that completely eliminate the adaptive immune repertoire, such as bone marrow transplantation (77-79).

## Methods

### Study Participants

Characteristics of study participants are provided in Supplementary Table S1. Participants were HIV-1-infected adults on suppressive ART initiated during chronic infection, with undetectable plasma HIV-1 RNA levels (<50 copies/ml) for ≥4 years. Additional inclusion criteria were CD4^+^ T cell counts >400 cells/μL and positive CMV serology. Peripheral blood samples (up to 180 mL) were collected at multiple time points (Figure. S2). For participants P9 and P10, leukapheresis was performed at a single time point.

### Isolation of CD8-depleted PBMCs

Peripheral blood mononuclear cells (PBMCs) were isolated and depleted of CD8^+^ T cells in a single step by density centrifugation using FicollPaque PLUS (GE Healthcare Life Sciences), SepMate tubes (Stemcell Technologies) and RosetteSep Human CD8 Depletion Cocktail (Stemcell Technologies) per the manufacturer’s instructions. For leukapheresis samples, total PBMCs were isolated by density centrifugation and viably frozen. PBMCs were thawed, rested overnight in RPMI with 10% FBS and, depleted of CD8^+^ T cells using a negative selection step (CD8 MicroBead Kit; LD columns, QuadMACS Separator, all from Miltenyi Biotec) per manufacturer’s instructions.

### Isolation of antigen-responding CD4+ T cells

To identify antigen-responding CD4^+^ T cells, CD8-depleted PBMCs were stimulated with antigen and co-stimulatory molecules (CD28+CD49d, BD Biosciences, 0.5 μg/mL) in the presence of CD40 blocking antibody to reduce CD40L internalization (Miltenyi, 1μg/mL) 10 μM enfuvirtide (T20) to prevent new infection events. We stimulated 10-100 million PBMCs with either lysates of CMV-infected fibroblasts (Virusys,10 μg/mL) or overlapping Gag 15mer peptides (HIV-1 Gag peptide pool, JPT Peptides, 1 μg/peptide/mL). For each stimulationaliquots of PBMCs (5-15 million cells) were cultured with no antigens (co-stimulation only control) or with anti-CD3/anti-CD28 antibodies bound to magnetics beads (25 μl/10^6^ cells, Thermo Fisher) (positive, nonspecific control). Cells were stimulated for 18 hours, ,washed and incubated with FcgR block (BD Pharmingen) at 25oC for 10 minutes before staining, for 30 min on ice with an APC-labelled antibody to CD3 (Biolegend; clone UCHT1), phycoerythrin (PE)-Cy7-labelled antibody to CD4 (Biolegend; clone RPA-T4), BV421-labelled antibody to CD45RO (Biolegend; clone UCHL1), PE-labelled antibody to CD154 (Biolegend; clone 24-31), FITC-labelled antibody to CD69 (Biolegend; clone FN50) and PE-Cy5-labelled antibodies to CD14 (ThermoFisher; clone 61D3), CD16 (Biolegend; clone 3G8) and CD20 (Biolegend; clone 2H7). Dead cells were excluded using propidium iodide. Cells stained with single fluorophore-labelled antibodies, co-stimulation only controls and positive controls were used to set sorting gates. Cells were sorted using either the Beckman Coulter MoFlo Legacy or XDP cell sorters. A representative gating strategy and sorting logic is provided in Figure 1 and S1A. To test non-specific activation of cells upon stimulation with CMV and Gag antigens, we stimulated PBMCs from 3 healthy donors negative for HIV-1 and CMV (Supplementary Figure S1C).

### Extraction of genomic DNA

Sorted cells were pelleted and immediately lysed for genomic DNA (gDNA) extraction. If samples had more than 10^6^ cells, we used the QIAamp DNA Mini Kit (Qiagen), otherwise we used the Quick-DNA Microprep Kit (Zymo), following manufacturer’s instructions. NanoDrop 2000 and Qubit 3 Broad Range (ThermoFisher) were used to quantify gDNA concentrations.

### HIV-1 DNA single genome sequencing

gDNA was subjected to limiting dilution nested PCR using the Platinum Taq High Fidelity DNA Polymerase (ThermoFisher), as previously described (80). To maximize chances of identifying identical proviral sequences across participants, we used a database of 431 near full length proviral sequences (81) to identify a 1.5 kb amplicon covering a region that is the least affected by deletions. In addition, we introduced degenerate positions in the primer sequences to allow amplification of sequences containing mismatches caused by viral diversification or APOBECG/F-induced G-to-A mutations. Primers were designed to amplify the U5-*gag* region (HXB2 positions 584-1841). A second set of primers was used to amplify a region of the *envelope* gene (HXB2 positions 6980-8036). Outer PCRs were performed with the following cycles: 94°C for 2 minutes, 44 cycles of: 94°C for 30 seconds, 50°C for 30 seconds, 72°C for 2’, then 72°C for 3’ and hold at 4°C. Inner PCRs were performed with the following cycles: 94°C for 2’, 41 cycles of: 94°C for 30 seconds, 55°C for 30 seconds, 72°C for 2’, then 72°C for 3’ and hold at 4°C. PCR product from positive reactions was purified and sent for direct Sanger sequencing (Genewiz). Oligonucleotide sequences used for PCR and Sanger sequencing are provided in Supplementary Table S6.

### Whole genome amplification

Antigen-responding cells from follow-up samples were sorted into 96 well plates in small pools (range 15-500 cells) to achieve, 30% or fewer HIV-1 DNA positive wellsSorted pools were then subjected to whole genome amplification by multiple displacement amplification (MDA) using the Advance Single Cell Repli-G kit (Qiagen), following manufacturer’s protocol. Whole genome amplified samples were diluted 1:20 in 10mM TrisHCl pH8.0, quantified by Qubit 3 Broad Range (ThermoFisher) and used as templates for HIV-1 bulk PCR (same methods used for single genome sequencing). Sanger sequencing from positive PCR reactions allowed the selection of HIV-1 positive wells containing individual proviruses of interest.

### Integration site analysis

Integration site analysis was performed using ligation-mediated PCR. In brief, whole genome amplified DNA from pools of cells positive for proviruses of interest were fragmented by sonication.DNA adapters were ligated onto the DNA ends. PCR was carried out between the adaptor and LTR ends, and then sequences were determined from both ends of PCR products using the Illumina method as described (82, 83). Abundance of individual clones was measured by quantifying the numbers of positions of DNA linkers associated with each unique integration site sequence, which scores the numbers of DNA chains at the start of the procedure associated with each cell clone (the sonic abundance method (84)). Some samples were analyzed with the Lenti-X Integration site analysis kit (Takara Bio), using primers designed to amplify both LTR ends. Libraries were prepared to capture the integration sites at both 5’ and 3’ host-provirus junctions. When necessary, primers were modified to reflect sequence variation in the HIV LTRs within the participant studied. Oligonucleotide sequences used for integration site analysis are given in Supplementary Table S6. The code base for the bioinformatic pipeline used to process high-throughput sequencing data is available here (doi: 10.5281/zenodo.3885688). Cancer-related genes were determined as previously described(82, 83). Expression of genes in CD4^+^ T cells and in other tissues was assessed through the Human Cell Atlas (http://immunecellatlas.net/) and the GTEx Project portal (https://www.gtexportal.org/home/). Only integration sites recovered from more than one sample and/or confirmed with a second method were used in the final analysis (see Supplementary Table S3). To recover integration sites in *BACH2* and *STAT5B* genes from previous publications, we used the NCI Retrovirus Integration Database (85). Only data from studies *in vivo* were used for the analysis in Figure 3B.

### Near full length proviral sequencing

To recover proviral sequences from whole genome amplified DNA, we used three different methods based on the type of provirus. Given the average size of DNA fragments generated by whole genome amplification (~3000 nt), we used 5 overlapping nested PCR amplicons (~2000nt each) as previously described (44). Proviruses with potential large deletions (inferred by the failure of one or more of the overlapping PCRs and IPDA) were amplified with outer and nested primers as described by Hiener *et al (27)*. Due to large deletions encompassing one of the LTR regions, some proviruses could not be amplified with this method. To bypass this issue, we designed primers in the human genome flanking the integration site. These were paired with HIV-specific primers annealing to proviral regions outside of inferred deletions, and used for nested PCR amplification using Platinum Taq High Fidelity DNA Polymerase (ThermoFisher) with modifications of the single genome sequencing protocol described above. Outer PCRs were performed with the following parameters: 94°C for 2’, 44 cycles of: 94°C for 30 seconds, 50°C for 30 seconds, 72°C for X’, then 72°C for 3’ and hold at 4°C. Inner PCRs were performed with the following parameters: 94°C for 2’, 41 cycles of: 94°C for 30 seconds, 55°C for 30 seconds, 72°C for X’, then 72°C for 3’ and hold at 4°C. The duration of the elongation step (X’) was adjusted to the estimated maximum size of the amplicon (1’ every 1000nt). Primer details are provided in Supplementary Table S6.

### Intact proviral DNA assay

The intact proviral DNA assay (IPDA) was performed as previously described(81). Due to sequence variation, a custom Ψ-probe was designed for participant P2 (FAM-TGGCGTACTCACCAGG-MGBNFQ, Applied Biosystems) and a custom Ψ-forward primer was designed for participant P3 (CAGGACTCGGCTTGCTGAGC).

### Quantitative viral outgrowth assay (qVOA)

qVOAs were performed as previously described (51). MOLT-4/CCR5 cells were added on day 2 and culture supernatants were examined for the p24 viral capsid protein by ELISA (PerkinElmer) after 14, 21 and 28 days. Results were expressed as infectious units per million cells (IUPM) CD4^+^ T cells calculated using maximal likelihood as previously described (86). Supernatants positive for p24 were collected, spun at 400 G for 10 min to remove cells and debris, and stored at −80°C. Cell-free HIV-RNA was extracted as previously described (87) and used for cDNA synthesis using SuperscriptIV (Invitrogen) as per manufacturer’s instructions. Reverse transcription was conducted at 55°C for 50’ followed by inactivation at 85°C for 10’, and held at 4°C, and cDNA was treated with RNAseH (1unit) at 37°C for 20’. For cDNA synthesis we used two separate reactions for the 5’ and the 3’ halves of the HIV-1 genome, with the following gene-specific primers: SCO5R (AGCTCTTCGTCGCTGTCTCCGCTT) and PARO (TTTTTGAAGCACTCAAGGCAAG). Five overlapping PCR amplicons were used to reconstruct partial or near full genome sequences, as previously described (44, 88). Primers used for direct Sanger sequencing have been previously published (18).

### Bioinformatic analysis of HIV-1 sequences

Raw data from Sanger sequencing was analyzed in Geneious to resolve base call conflicts and eliminate sequences with poor quality or double peaks (more than one HIV-1 variant per PCR reaction). Sequence contigs were aligned with ClustalW(89), G > A hypermutants were identified with Hypermut 2.0 (https://www.hiv.lanl.gov/content/sequence/HYPERMUT/hypermut.html) and codon alignments were analyzed with GeneCutter (https://www.hiv.lanl.gov/content/sequence/GENE_CUTTER/cutter.html) to identify premature stop codons and frameshifts. To confirm the lack of genetic defects affecting infectivity, we used the Proviral Sequence Annotation & Intactness Test (Proseq-IT)(90). ElimDupes was used to identify and collapse identical sequences (https://www.hiv.lanl.gov/content/sequence/elimdupesv2/elimdupes.html). Neighbor-Joining trees were constructed in MEGA 7.0(91) with a subtype-specific HIV-1 consensus as the outgroup. Phylogenetic structure was tested by bootstrap analysis (1000 replicates). To estimate the oligoclonality of proviral populations, we used the Gini coefficient of inequality(92), calculated in RStudio with the *ineq* R package (https://cran.rproject.org/web/packages/ineq/index.html) and corrected for small samples(93). This measurement of sample dispersion provides an estimate of whether the proviral sequences within a sample are evenly distributed (values approaching 0) or dominated by groups of identical sequences (values approaching 1).

### TCRβ Immunosequencing

Genomic DNA was isolated from cell samples, quantified as described above and diluted in Tris-acetate-EDTA to a concentration of 10ng/uL (for a total of up to 1ug per sample). TCRβ sequencing data were generated using the ImmunoSEQ hsTCRB assay, version 4, survey mode (Adaptive Biotechnologies).

### TCRβ analysis

TCRβ sequencing data were analyzed with ImmunoSeq Analyzer 3.0 (Adaptive Biotechnologies). Diversity analyses were conducted on nucleotide sequences from productive VDJ rearrangements. The lower bound estimate of diversity was calculated with the improved Chao index (94). Estimate of TCR clonality was based on the Gini coefficient (as described above) calculated in R with the *Immunarch* package (https://cloud.r-project.org/package=immunarch). Differential abundance analysis of VDJ rearrangements between populations of sorted cells was conducted in ImmunoSeq Analyzer 3.0, as previously described (95). Rearrangements with different nucleotide sequences encoding for the same CDR3β amino acid sequences were considered as distinct clonotypes. Identification and tallying of degenerate CDR3β sequences were performed in Windows Excel. We used GLIPH2 (55) to cluster TCRs based on predicted binding to peptide-loaded MHC. GLIPH2 pipeline was applied on TCR data sets obtained from individual participants and merged sets from all participants. For clustering based on global similarity, member CDR3s need to be of the same length, and differ at the same position. Amino acids were interchangeable only if had a positive score in BLOSUM-62 matrix (55). Analysis and visualization of significant CDR3β convergence groups were performed in RStudio. Additional details of the four TCR clusters shown in Figure 4I are provided in Supplementary Table S4. Visualization of CDR3β networks and sequences logos were conducted with Gephi (https://gephi.org/) and Weblogo (96), respectively. HLA haplotypes of the participants are shown in Supplementary Table S7.

### T cell subset analysis

A fraction of CD8-depleted PBMCs, isolated as described above, were dedicated to sort CD4^+^ T cell subsets. We incubated cells with FcgR block (BD Pharmingen) at room temperature for 10 minutes. Cells were stained, with a 30 minutes incubation on ice, with a APC-labelled antibody to CD3 (Biolegend; clone UCHT1), phycoerythrin (PE)-Cy7-labelled antibody to CD4 (Biolegend; clone RPA-T4), BV421-labelled antibody to CD45RA (BD Biosciences; clone HI100), PE-labelled antibody to CCR7 (Biolegend; clone G043H7) and PE-Cy5-labelled antibodies to CD14 (Thermo Fisher; clone 61D3), CD16 (Biolegend; clone 3G8) and CD20 (Biolegend; clone 2H7). Dead cells were excluded using propidium iodide. Cells stained with single fluorophore-labelled antibodies were used to set sorting gates. A representative gating strategy and is provided in Figure S10A. CD45RA expression was used to distinguish Naïve-like and terminally differentiated cells from memory cells. Central memory cells were distinguished from effector memory cells by the expression of CCR7. Naïve-like (Na), Central Memory (CM), Effector Memory (EM) and Effector Memory CD45RA positive (EMRA) subsets were sorted using either the Beckman Coulter MoFlo Legacy or XDP cell sorters.

### Quantification of specific proviruses and VDJ rearrangements

See supplementary methods.

### Assignment of TCR-provirus pairs from Antigen-reactive clonotypes

See supplementary methods.

### Total body size estimates and likelihood of homeostatic proliferation

See supplementary methods.

### Quantification and Statistical analysis

Descriptive statistics, tests for normality, spearman correlation, 2-tailed Student’s t-test and one-way ANOVA tests were used to determine statistical significance using GraphPad Prism v8.0. A P value less than 0.05 was considered significant, unless otherwise stated.

### Study Approval

The Johns Hopkins Institutional Review Board and the UCSF Committee on Human Research approved this study. All participants provided written consent before enrolment.

## Data and Software availability

HIV-1 sequences are available on GenBank (accession numbers MW255985-MW256411, MW262767-MW262790 and MW309883-MW309909). Integration site sequencing data can be found on the SRA database at this link: https://www.ncbi.nlm.nih.gov/sra/PRJNA637643. The code base for the bioinformatic pipeline used to process high-throughput integration site sequencing data is available here (doi: 10.5281/zenodo.3885688). Integration site data has also been submitted to the NCI Retrovirus Integration Database (https://rid.ncifcrf.gov). TCRβ sequencing data can be accessed through the ImmuneAccess database (doi: 10.21417/FRS2020JCI, url: clients.adaptivebiotech.com/pub/simonetti-2020-jci).

## Author contributions

F.R.S. and R.F.S. conceptualized the study. F.R.S., G.S., J.D., K.R., S.A.B., K.K., J.W., J.L., performed experiments. H.Z. and J.B.M. performed cell sorting. K.M., H.R, C.N., S.E. and F.D.B. conducted integration site analysis. A.L.H. provided mathematical model of homeostatic proliferation. F.R.S. analyzed the data and generated figures. S.A.B., R.H. and S.G.D. provided participant samples. F.R.S. and R.F.S. wrote the manuscript with input from all authors.

## Acknowledgements

We thank the participants who volunteered in this study and their families. We thank Guido Massaccesi and Andrea L. Cox for providing PBMCs from donors with known cell-mediated responses to CMV. We thank Srona Sengupta, Annie A. Antar, Janelle Montagne and H. Benjamin Larman for discussions leading to this work, and Monica Sullivan for the precious administrative support. This work was supported by the NIH Martin Delaney, Beat-HIV (UM1 AI126620) and DARE (UM1 AI12661) Collaboratories, by the Howard Hughes Medical Institute and the Bill and Melinda Gates Foundation (OPP1115715). This work was also supported by NIH grants DP5OD019851, U19-AI117950, R01AI129661, R01CA241762, the Penn Center for AIDS Research (P30AI045008) and the PennCHOP Microbiome Program.

