## Supplementary Materials for "Antigen-driven clonal selection shapes the persistence of HIV-1 infected CD4^+^ T cells *in vivo*"

**Supplementary Results.** Characterization of proviral sequences in antigen-responding clones.

**Supplementary Methods.**

**Supplementary Figure S1.** Accessing antigen-responding CD4+ T cells.

**Supplementary Figure S2.** Longitudinal sampling schema and experiments performed at each time point.

**Supplementary Figure S3.** Validation experiment of whole genome amplification on sorted JLat cells

**Supplementary Figure S4.** Summary of identical HIV-1 DNA single genome sequences recovered from sorted cells.

**Supplementary Figure S5.** Proviral structures with large deletions and aberrant provirus/host junctions.

**Supplementary Figure S6.** Intact proviral DNA assay analysis of individual proviruses.

**Supplementary Figure S7.** Quantitative viral outgrowth assay data from sorted CD4+ T cells.

**Supplementary Figure S8.** Additional analyses on TCRβ sequences.

**Supplementary Figure S9.** Pairing of TCRβ sequences and proviruses belonging to the same clonotype.

**Supplementary Figure S10.** Contribution of CD4+ T cell subsets to antigen-responsive clonotypes carrying HIV proviruses.

**Supplementary Table S1.** Participant characteristics.

**Supplementary Table S2.** Single genome sequences and oligoclonality of proviral populations from sorted CD4+ T cells.

**Supplementary Table S3.** Integration site analysis of HIV-1-infected antigen-responding clones.

**Supplementary Table S4.** Characteristics of TCRβ sequences in clusters shown in Figure 4.

**Supplementary Table S5.** Likelihoods that clones would reach the observed sizes under a model of homeostatic proliferation.

**Supplementary Table S6.** Primers and probes used in this study (xml file).

**Supplementary Table S7.** High resolution HLA genotyping of the study participants.

### Supplementary Results

#### Characterization of proviral sequences in antigen-responding clones.

To investigate whether clones of HIV-1-infected, Ag-responding cells carried intact or defective proviruses, we recovered the partial or full-length sequences of the proviruses for which we identified integration sites (Figures 3A and C). As expected, most genomes (13/22) were defective due to deletions that we either mapped (8/22) or inferred (5/22) due to failure of PCR amplification or lack of signal from one amplicon in the Intact Proviral DNA assay (IPDA, see below). Of the remaining defective proviruses, seven showed G-to-A hypermutation and one had a 93-nucleotide deletion affecting the dimerization, major splicing donor, and packaging signal stem loops (P5, integrated in *DELEC1*). Only one provirus had no defects that would abrogate infectivity (found in participant P3, integrated in *FXBO22*).

In some cases, primers commonly used for near full-length proviral sequencing (27, 97, 98) failed due to deletions. Therefore, we used the integration site to design primers in host DNA flanking the host-proviral junctions. This allowed recovery of near full genome sequences from 5 proviruses (integrated in *ST6GALNAC3*, *KCNC2* and 3 proviruses in *BACH2*, respectively) with unusual proviral structures (Figure S5). These proviruses had large deletions (range 3749-7335 nucleotides) almost completely encompassing one of the LTRs (Figures 3A and S5). For two of these proviruses, the deletion extended outside the HIV-1 genome, and the 5-nucleotide duplication typically left by the integration process (99) was absent, suggesting that these proviruses might be the result of aberrant integration (100, 101). These observations reflect the striking heterogeneity of the proviral landscape, which is challenging to analyze without PCR bias. Two of these proviruses (integrated in *ST6GALNAC3* and outside of *KCNC2*) belonged to the most abundant infected clones in the study but would have been missed with standard full-genome sequencing.

We also analyzed whole genome amplified DNA using the IPDA, a novel droplet digital PCR assay that uses two strategically placed amplicons to distinguish intact from defective proviruses (81). The IPDA allowed assessment of the intactness of these clonally expanded proviruses in case of incomplete recovery of genome sequences (Figure 3A). In addition, for 17 completely sequenced proviruses, the IPDA signal pattern showed 100% agreement with patterns expected based on the sequencing data (Figure S6).

### Supplementary Methods

**Quantification of specific proviruses and VDJ rearrangements.** To quantify proviruses of interest together with the VDJ rearrangement of their cognate clonotypes, we used the droplet digital PCR platform QX200 (Bio-Rad). In order to prevent non-specific amplification of unrelated proviral sequences, we designed primers across the host-U3 or U5-host junctions. In addition, the fluorescently labelled probe was designed to anneal across the site of HIV-1 integration. In case of proviruses integrated in regions with repetitive sequences or with a GC content unsuitable for primer design, we designed primers and probes across large internal deletions that were unique to the provirus of interest. Similarly, for primers and probes aimed to quantify specific VDJ regions, we placed the forward and reverse primers on the V and J regions, respectively, and the probe was designed to span the junctions between the V-D-J regions, in order to exploit N and P nucleotides and small deletions randomly occurring during the recombination process. PCR reactions were run with the following parameters: 95°C for 10’, 95°C for 30s, 56°C for 2’ (steps 2 to 3 for 44 cycles), 98°C for 10’, hold at 4°C (temperature change rate 2°C/s). Primers and probe had a final concentration of 900nM and 250nM, respectively. The specificity of each set was tested against a panel of whole genome amplified DNA samples with and without the $CD4^+$ T cell clones of interest and a sample of $CD4^+$ T cell derived gDNA from healthy donors. Primers are probes are shown in Supplementary Table S6. We performed parallel quantification of RPP30 as previously described (81) to calculate cell equivalents and normalized copies of proviral integration sites and VDJ rearrangements.

**Assignment of TCR-provirus pairs from Antigen-reactive clonotypes.** To pair the TCRβ of an antigen-responsive CD4+ cells and the provirus integrated in the same clonotype, we leveraged TCR diversity and combinatorial statistics previously used to confidently pair TCR α and β chains (56). Briefly, a fixed number of antigen-reactive cells are randomly allocated to each well on a 96-well plate and subjected to WGA as described above. TCRβ immunosequencing is performed on wells sharing the same proviral sequence and integration site and TCRβ sequences recurring in all wells are considered as candidate for pairing experiments. Subsequently, the whole plate is screened by duplex ddPCR with assays design to specifically detect both the integration site and the candidate TCRβ sequence. Given that immune repertoire is highly diverse, even within antigen-responsive cells, the probability that a specific TCRβ sequence and a provirus will occupy exactly the same sets of wells is miniscule, so a pair of TCRβ and proviral sequences that uniquely share a set of wells can be inferred to have come from the same clone. For each potential TCRβ/provirus pair, we computed a *P* value for the observed number of shared wells, implementing the calculations described by Howie *et al.* (56) shown in supplementary Figure S9A and B.

**Total body size estimates and likelihood of homeostatic proliferation.** To estimate the total body size of HIV-1-infected CMV-responding clones, we used the frequency of infected cells carrying the relevant provirus among all CMV-responding $CD4^+$ T cells, as quantified with the provirus-specific ddPCR assays. We then used the percentage of CMV-responding $CD4^+$ T cells upon antigen stimulation to extrapolate the frequency of the relevant clone among all $CD4^+$ T cells. Total body cell number ($2^{11}$) was estimated based on the average $CD4^+$ T cell counts in individuals on long-term ART (17, 18, 27) (800 cell/µL), the fraction of cells in circulation (102) (0.02) and the average total blood volume (5L). The number of divisions needed to reach the observed clone sizes is calculated with the $Log_2$ of the total body size of each infected clone (Supplementary Table S5). Of note, this assumes no cell death (a clone can only divide). We then calculated the likelihood a clone could reach this size if the entire population of $CD4^+$ T cells was maintained by a constant, balanced, process of division and death (only random events lead clones to grow or shrink). We applied a range of proliferation rates (10-1000 divisions/year) to represent cell subsets with different proliferative potential (i.e. naïve vs resting memory vs activated effector cells). Calculations were conducted with two scenarios, one with the expected reservoir decay (4)(5) (44 months) and one with no net decay (103) (defective or total HIV-DNA). The death rate ($d$) was calculated so that the difference between the proliferation rate ($p$) and the death rate was equal to the observed net decay rate ($\delta = p - d$). The probability of observing a clone of size $X$ or larger with these turnover parameters was calculated using the formula

$$\bar{F}(X;t,p,d) = \begin{cases} 1 & X = 0 \\ (1-\alpha)\beta^{X-1} & X > 0 \end{cases}$$

With

$$\alpha = \frac{de^{\delta t} - d}{pe^{\delta t} - d},$$
$$\beta = \frac{pe^{\delta t} - p}{pe^{\delta t} - d}$$

which is the complementary cumulative distribution function for the solution of a homogeneous birth-death process starting with a single cell at time zero (104). We take time zero to be the time for viral suppression to < 50 copies per mL, which means we assume that proliferation predominantly occurs after viral suppression. If there are $N$ unique clones in the body, then the likelihood of observing at least one clone that is at least size $X$ is

$$\mathcal{L}(X;t,p,d) = 1 - (1 - \bar{F}(X;t,p,d))^N \approx N\bar{F}(X;t,p,d)$$

where the approximation holds as long as $\bar{F}(X;t,p,d) \ll 1$ . We assume that at the time of viral suppression, all clones are unique, so $N$ is equivalent to the estimate of the total number of latently infected cells at that time. Parameters and likelihoods for each clone are shown in Supplementary Table S5.

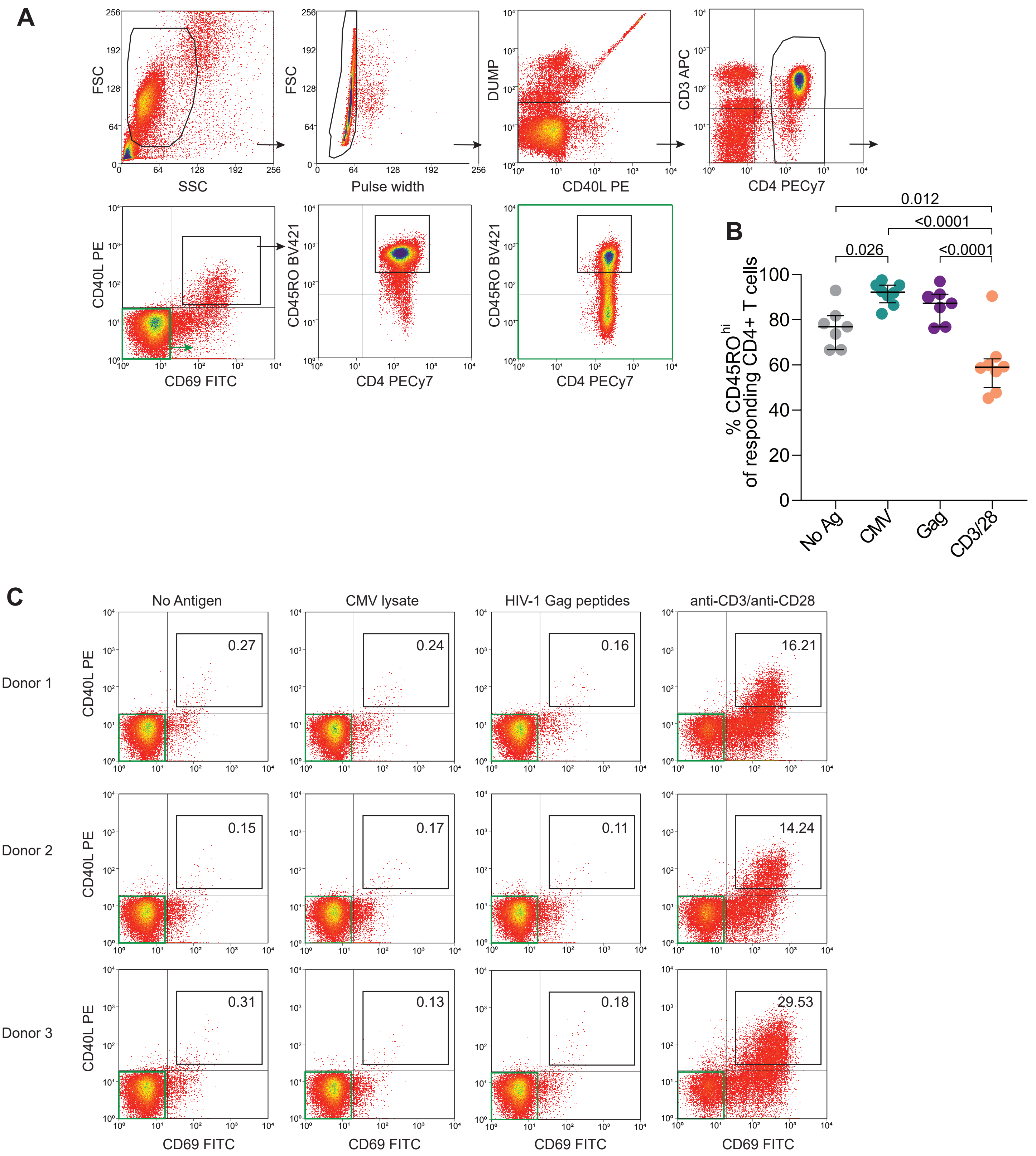


**Supplementary Figure S1. Accessing antigen-responding CD4⁺ T cells.**
(**A**) Gating strategy used to isolate CD4⁺ T cells responding to stimulation. Flow data from a representative stimulation of CD8-depleted PBMC with CMV lysate is shown. Non-responding cells gated on CD45RO expression are highlighted in green. (**B**) Fraction of CD4⁺ T cells responding to the indicated stimulations that are CD45RO$^{hi}$. Statistical significance was determined by one-way ANOVA test. (**C**) Frequency of CD4L and CD69 double positive cells among CD4⁺ T cells in response to stimulation of PBMCs from three donors naive for CMV and HIV-1 infection. Numbers in gates indicate percentage of double positive events.

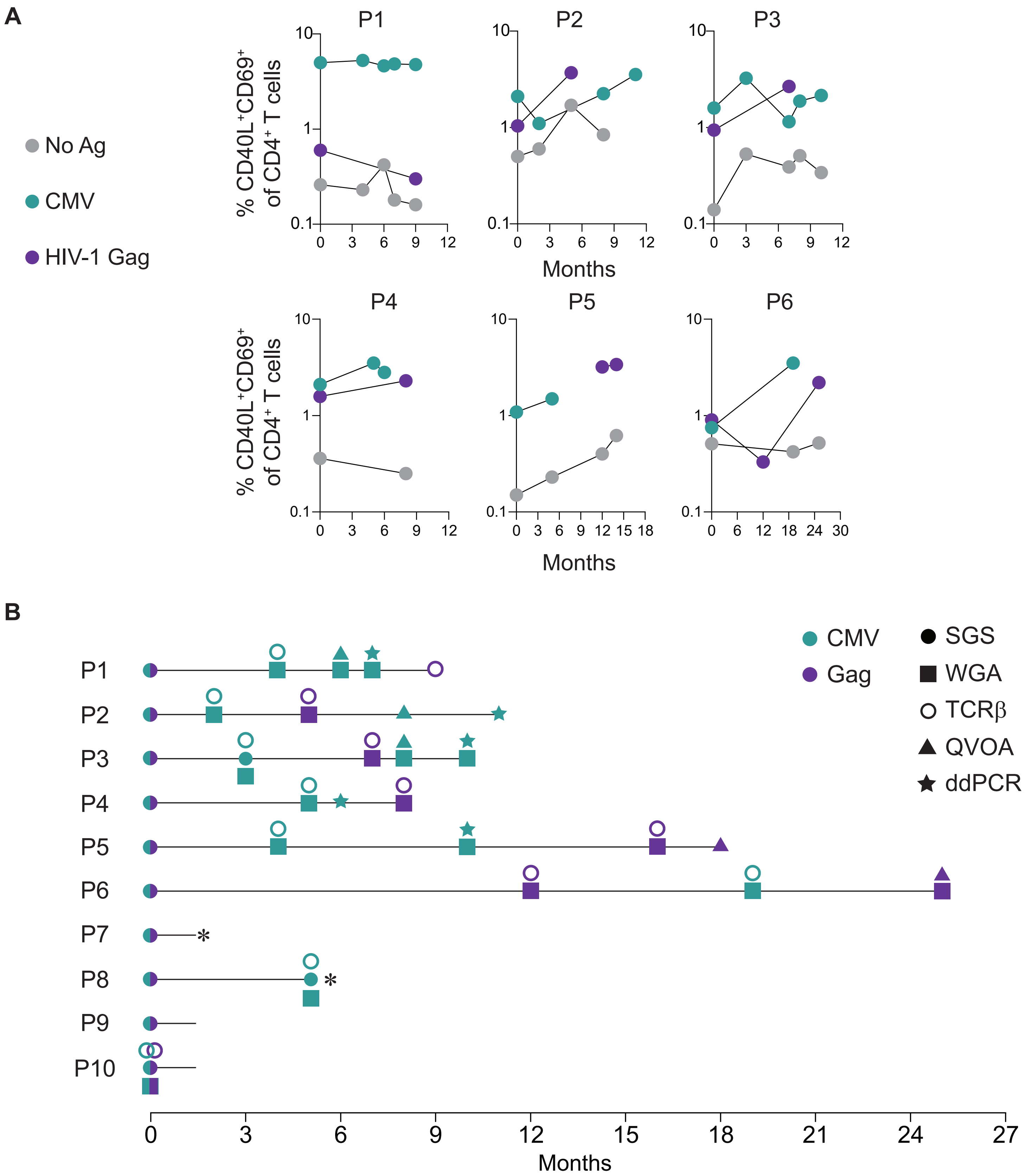


**Supplementary Figure S2. Longitudinal sampling schema and experiments performed at each time point.** (**A**) Frequencies of cells responding to the indicated stimulations in longitudinal samples from 6 participants with at least 3 time points available. (**B**) Each horizontal line represents one participant. Peripheral blood was collected from longitudinal samples at the indicated time points. Experiments performed on sorted cells are represented as described in the symbol legend. Participants P7 and P8 were lost to follow up (asterisk) due to ART interruption and fatal acute coronary syndrome, respectively.

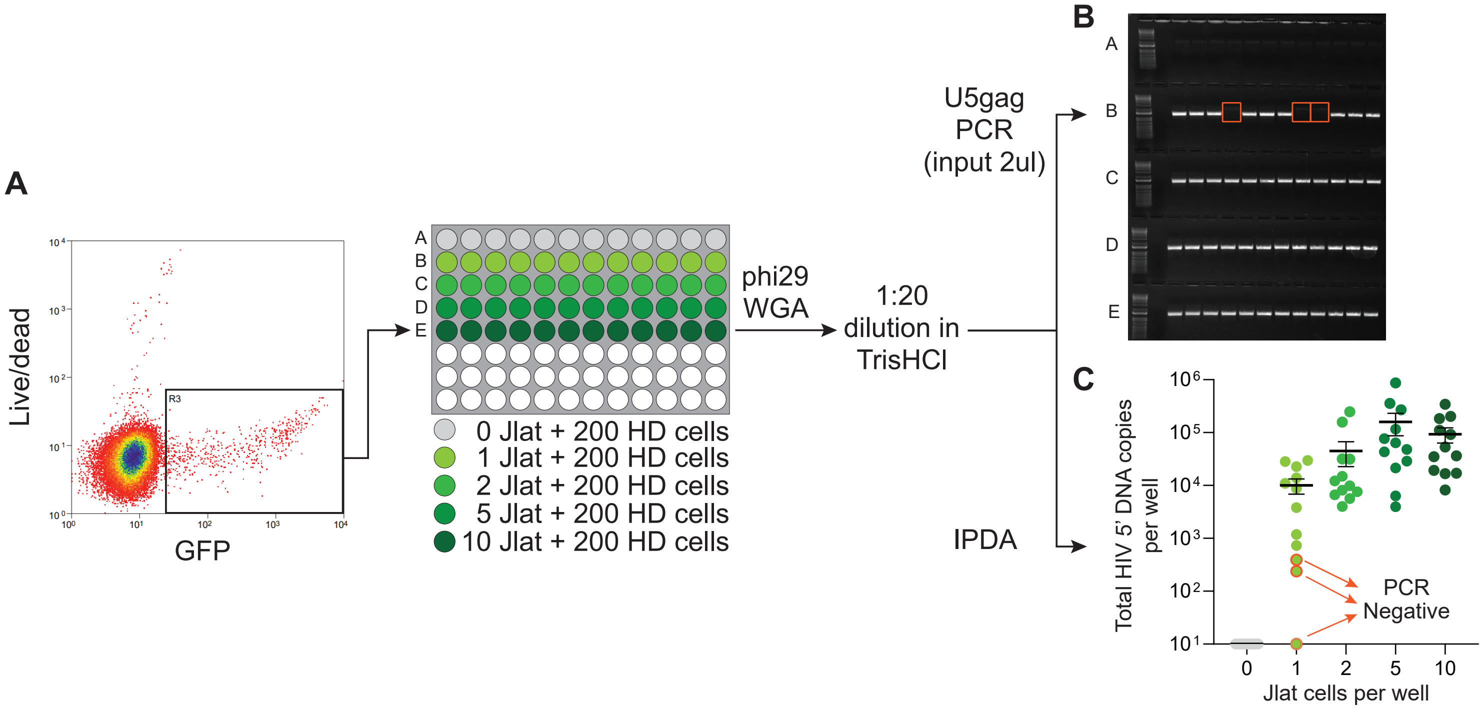


**Supplementary Figure S3. Validation experiment of whole genome amplification on sorted JLat cells.** (**A**) J-Lat cells (clone 10.6) were sorted into wells with 200 CD4⁺ T cells from a healthy donor. Cell pools were subjected to whole genome amplification and a diluted aliquot was used for downstream experiments. (**B**) Electrophoresis gel showing amplification of u5-gag PCR. Reactions lacking expected amplification are highlighted in orange boxes. (**C**) Quantification of 5' HIV genome by IPDA from whole genome amplified wells. Each dot represents the estimate of total HIV genome copies per well. Horizontal bars show mean and standard error of the mean.

A

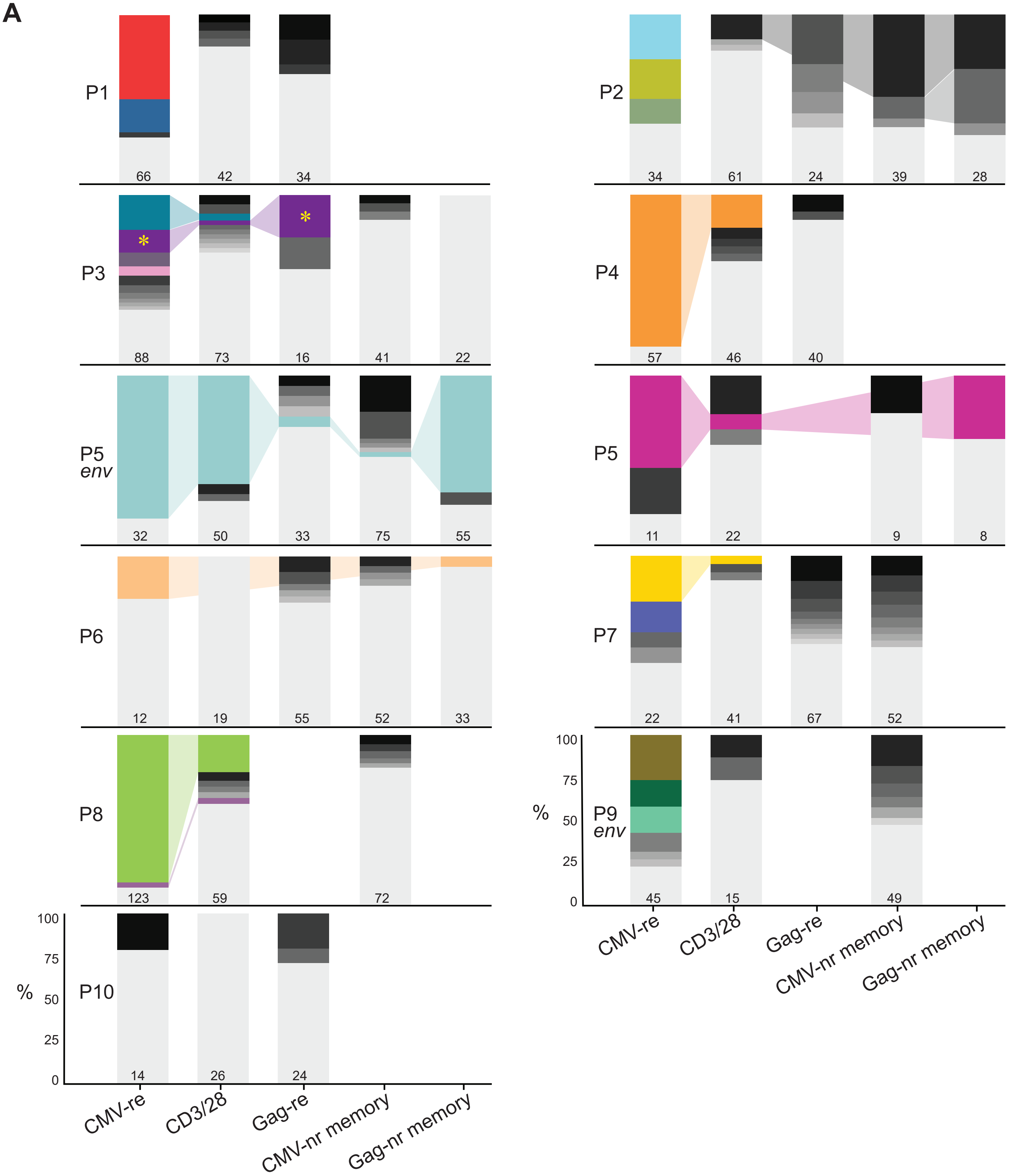


**Supplementary Figure S4. Summary of identical HIV-1 DNA single genome sequences recovered from sorted cells.**

(**A**) Stacked bar graphs represent proviral variants ranked by frequency. Variants identified at least twice are shown in shades of grey; clones of interest are color-coded as in main Figure 2C and D; singletons are shown in light grey at the bottom of the stack; potential clones found across more than one cell fraction are connected with shaded colors; the yellow star symbol highlights a case of a proviral sequences enriched both upon stimulation with CMV and Gag antigens.

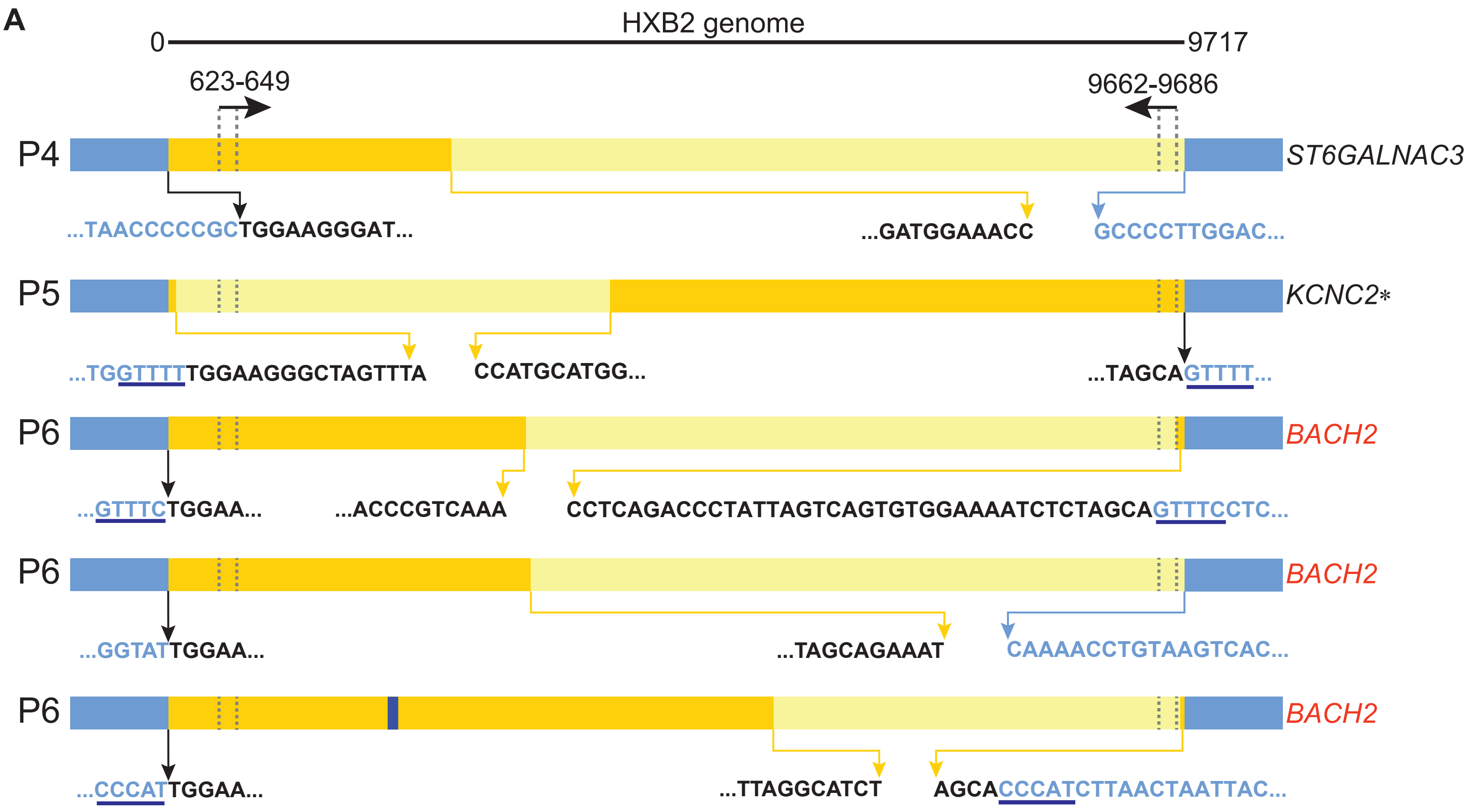


**Supplementary Figure S5. Proviral structures with large deletions and aberrant provirus/host junctions.** (**A**) Horizontal bars show proviral structures as displayed in Figure 3A. Black horizontal arrows and dashed vertical lines show the location of commonly used outer PCR primers (position numbers relative to HXB2 reference). Vertical arrows show the sequence flanking deletions and host-provirus junctions. Nucleotides belonging to the flanking host sequence are colored in blue. Five-nucleotide duplications, result of integrase-mediated insertion, are underscored in dark blue.

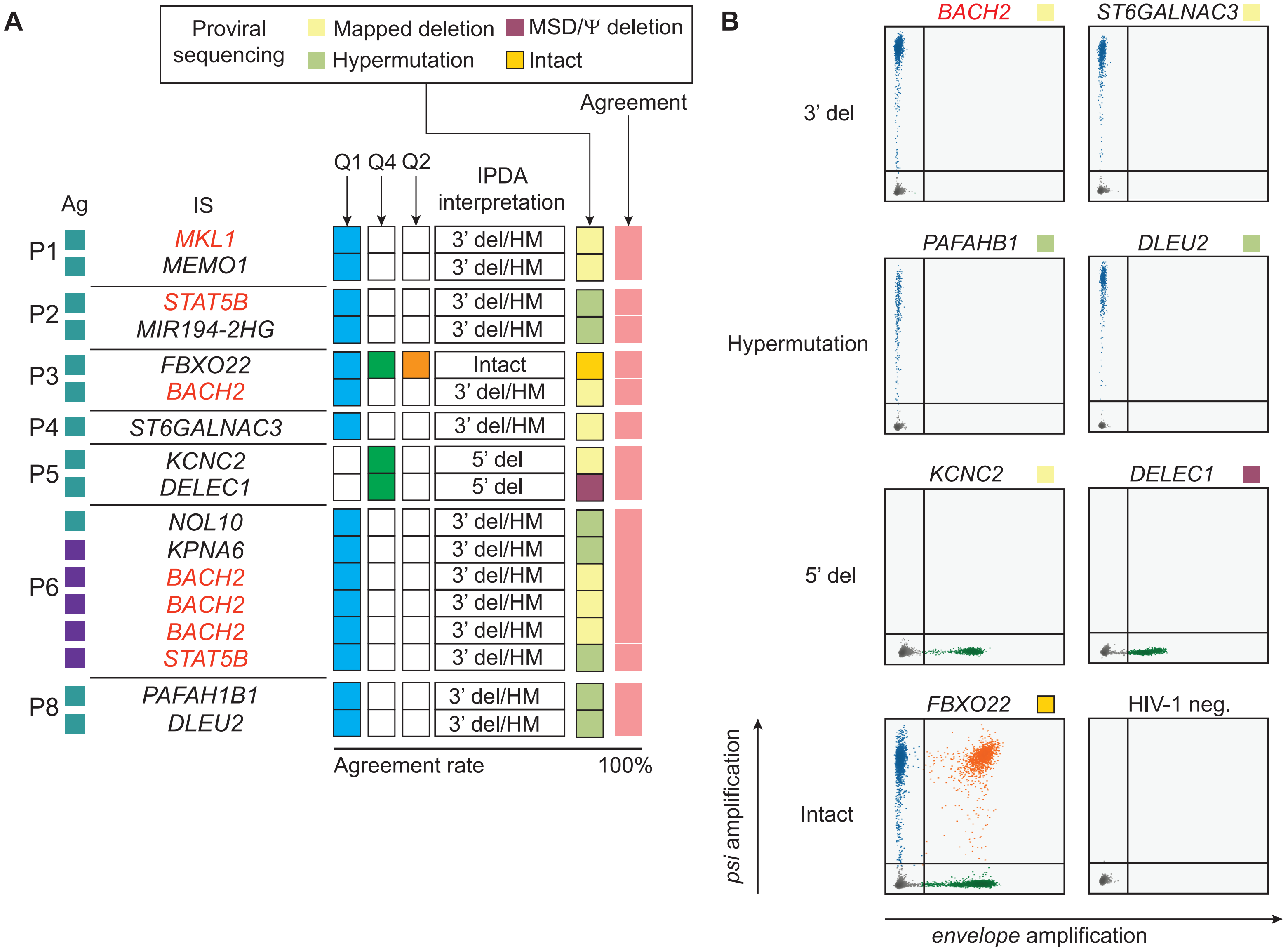


**Supplementary FIgure S6. Analysis of Intact proviral DNA assay performed on individual proviruses.**
(**A**) Summary of IPDA results from whole genome amplified genomic DNA samples containing individual proviruses (related to Figure 3A). Interpretation of IPDA signal is compared to proviral sequences, with features color-coded according to the boxed legend. Agreement between IPDA and proviral sequencing is highlighted in pink. (**C**) Examples of two-dimension ddPCR plots from the indicated proviruses with known 3’ deletions or hypermutation, 5’ deletions or small major splice donor/packaging signal defects, and one intact provirus.

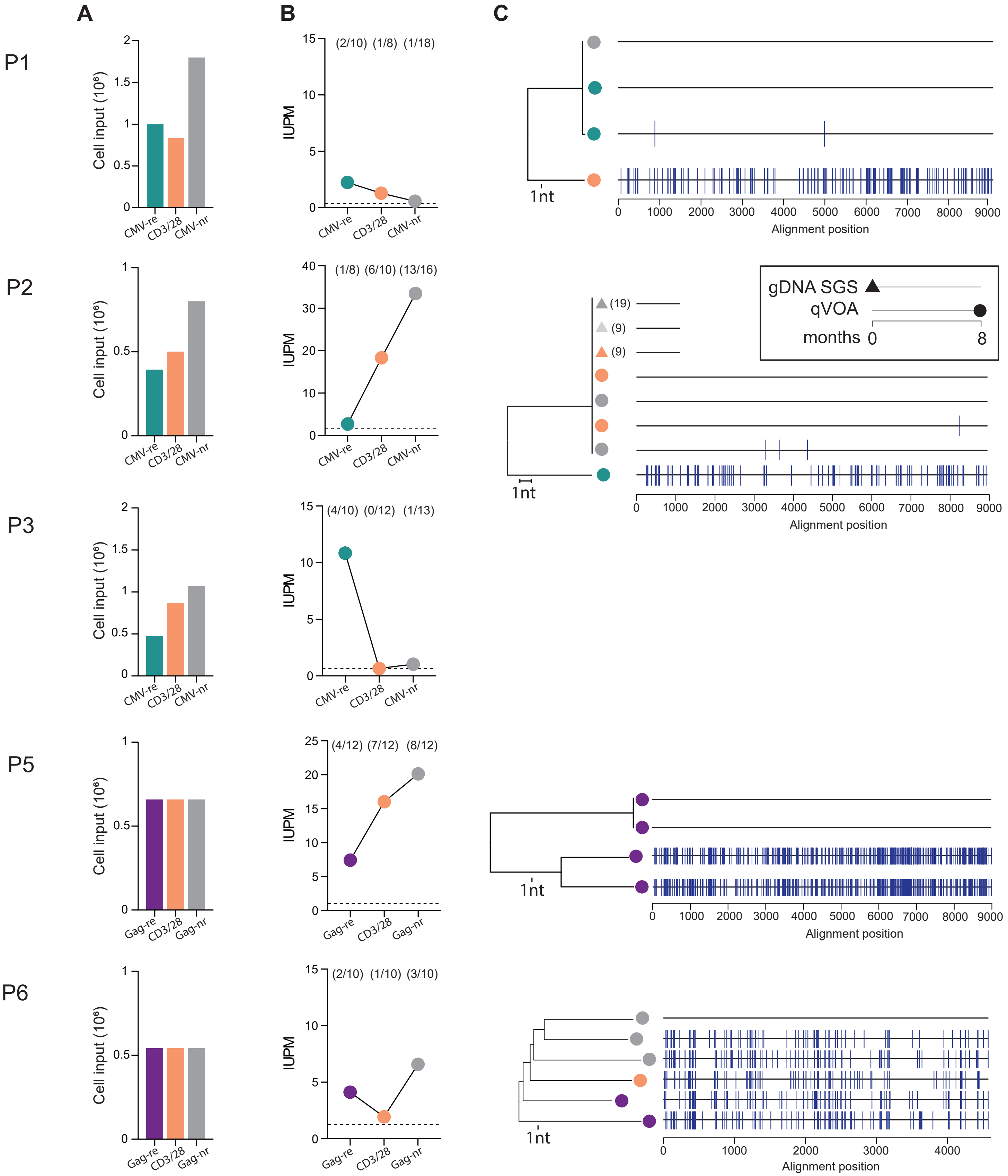


**Supplementary FIgure S7. Quantitative viral outgrowth assay data from sorted CD4⁺ T cells.**
(**A**) CD8-depleted PBMCs were stimulated as in the previous experiments (P1, P2, P3 with CMV, P4 and P5 with Gag, respectively) and the sorted antigen-responding cells were immediately used as input for the quantitative viral outgrowth assay (qVOA). The number of cells used as input for qVOA experiments for each participant are displayed in bar graphs. (**B**) Frequency of cells with inducible replication-competent proviruses in IUPM from each sorted cell fraction. Numbers in parentheses indicate the number of p24-positive wells out of all replicates performed for each condition. (**C**) Neighbor-Joining (NJ) trees of sequences from p24-positive wells and highlighter plots showing mismatches relative to the top sequence in the tree. In participants P1, P5 and P6, the NJ tree was build based with the same sequences used for the highlighter plot. In P2, NJ tree is based on u5-gag sequences. Source and time point of sequences are described in the insert. Sequence analysis for participant P3 is shown in Figure 3E.

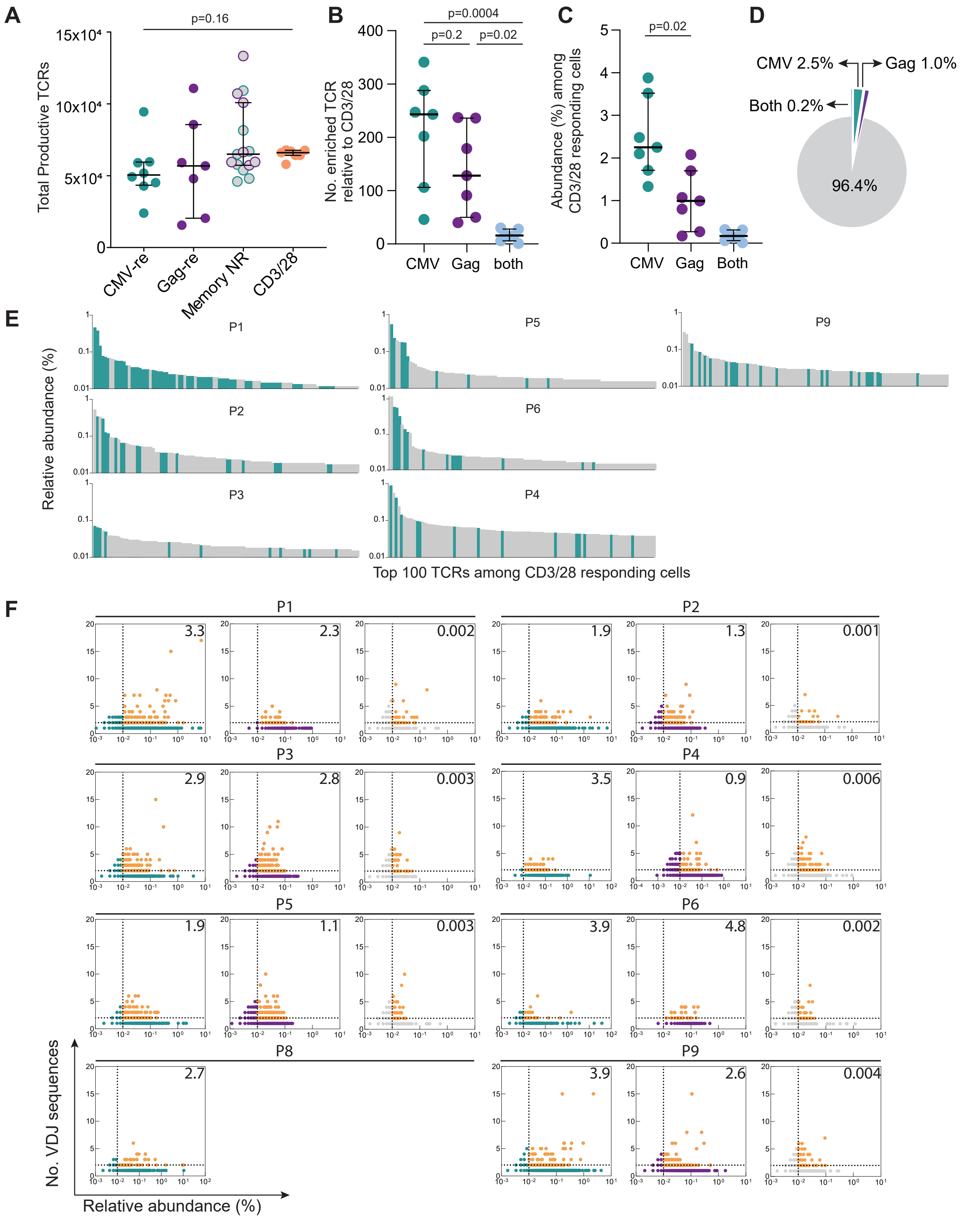


**Supplementary Figure S8. Additional analyses on TCR sequences related to Figure 4.**
(**A**) Total productive TCR sequences recovered from sorted cells. Grey circles indicate CD45ROhi cells non responding to either CMV or Gag stimulation (teal and purple border, respectively) (**B**) Number of TCR sequences from CD3/CD28-responding cells with a significantly higher abundance in CMV- or Gag-responding cells, or both. Horizontal bars show median and interquartile values. Statistical significance was determined using a one-way ANOVA. (**C**) Relative abundance (%) of TCR sequences enriched in antigen-responding cells among cells activated with a nonspecific stimulation. (**D**) CMV- and Gag-responding cells represent only a small fraction of all CD3/CD28-responding cells (mean values) (**E**) Relative abundance (%) of the top 100 clones from CD3/CD28-responding cells. Clones significantly enriched in CMV-responding cells are highlighted in teal. (**F**) Unique and degenerate CDR3β sequences were plotted against their relative productive abundance. Degenerate CDR3β sequences with a sum productive abundance higher than 0.01% are highlighted in orange (their cumulative abundance is indicated in the right upper quadrant).

A

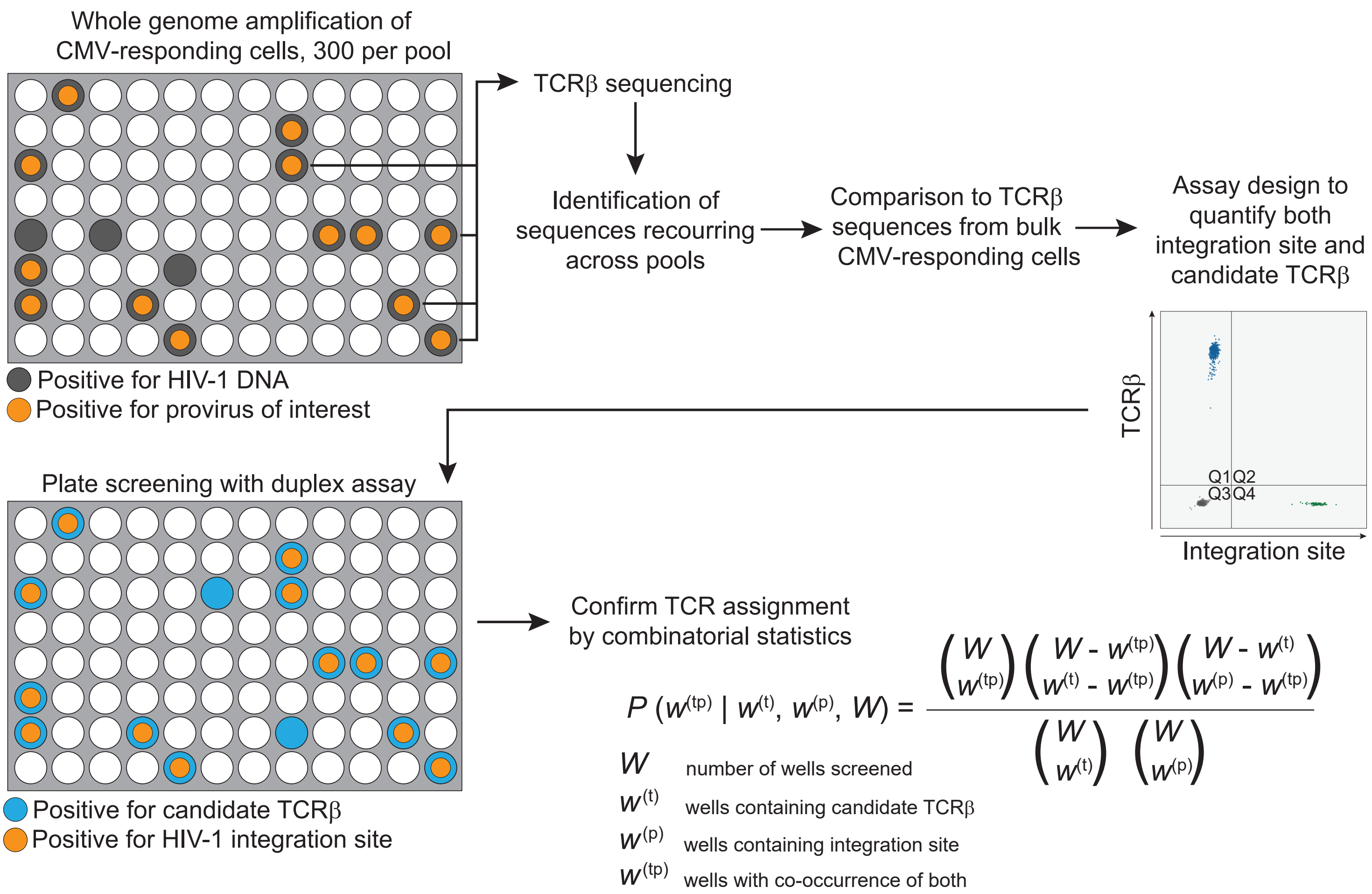


B

| Participant ID | TCRβ | Provirus | Cells per well | $W$ | $w^{(t)}$ | $w^{(p)}$ | $w^{(tp)}$ | $P$ | Provirus:TCR ratio (WGA) | Provirus:TCR ratio (ddPCR) |
|---|---|---|---|---|---|---|---|---|---|---|
| P1 | CASIGSSAFF | MKL1 | 100 | 92 | 41 | 7 | 7 | **$2.5 \times 10^{-3}$** | 0.17 | 0.14 |
| | | | 300 | 90 | 75 | 15 | 15 | **0.049** | 0.20 | |
| P3 | CASSYSTGITEAFF | FBXO22 | 200 | 96 | 16 | 3 | 3 | **$3.9 \times 10^{-3}$** | 0.19 | 0.15 |
| | | | 400 | 93 | 23 | 6 | 5 | **$3.0 \times 10^{-3}$** | 0.22 | |
| P4 | CASSLLTAATNEKLFF | ST6GALNAC3 | 20 | 90 | 5 | 5 | 5 | **$2.2 \times 10^{-8}$** | 1.00 | 0.72 |
| | | | 60 | 96 | 10 | 5 | 5 | **$4.0 \times 10^{-6}$** | 0.50 | |
| P5 | CASSLSGTENSPLHF | KCNC2 | 15 | 86 | 15 | 12 | 8 | **$4.0 \times 10^{-5}$** | 0.53 | 0.93 |
| | | | 30 | 92 | 37 | 42 | 33 | **$1.0 \times 10^{-10}$** | 0.89 | |
| | CSARATTGELFF | DELEC1 | 300 | 96 | 14 | 12 | 12 | **$1.4 \times 10^{-13}$** | 0.86 | 0.92 |
| P6 | CSVARQGAVYEQFF | BACH2.c1 | 100 | 96 | 7 | 7 | 7 | **$8.3 \times 10^{-11}$** | 1.00 | na |
| | CASSITGVGGQPQHF | BACH2.c2 | 100 | 95 | 6 | 5 | 5 | **$1.0 \times 10^{-7}$** | 0.83 | na |
| | CSVPGLGAEAFF | BACH2.c3 | 100 | 95 | 4 | 4 | 4 | **$3.1 \times 10^{-7}$** | 1.00 | na |

**Supplementary Figure S9. Pairing of TCRβ sequences and proviruses belonging to the same clonotype.** (**A**) Experimental approach to pair TCR sequences and proviruses belonging to the same antigen-responding clone. A representative example from participant P5 is shown (CSARATTGELFF-*DELEC1* pair). Whole genome amplified cell pools sharing identical proviral sequences and integration sites are subjected to TCR sequencing. Shared TCR sequences are identified and compared to TCR repertoire from bulk antigen-responding cells. If a unique TCR sequence is identified, the pair is confirmed by duplex ddPCR assays designed to amplify both the TCR and the integration site (see methods for details). For each potential TCRβ-provirus pair, we screened one or more plates with pools of antigen-reactives cells subjected to WGA and recorded the pattern of occurrence of wells positive for TCR and integration sites. We then computed a P value for the observed number of shared wells, implementing the calculations described by Howie *et al.*(56) (**B**) Confirmed TCRβ-provirus pairs and P values supporting that their pattern of co-occurrence is driven by chance. Of note, the provirus:TCR ratios inferred by the occurrence pattern correlate with the ratios calculated by ddPCR in antigen-responding cells (Figure 5C).

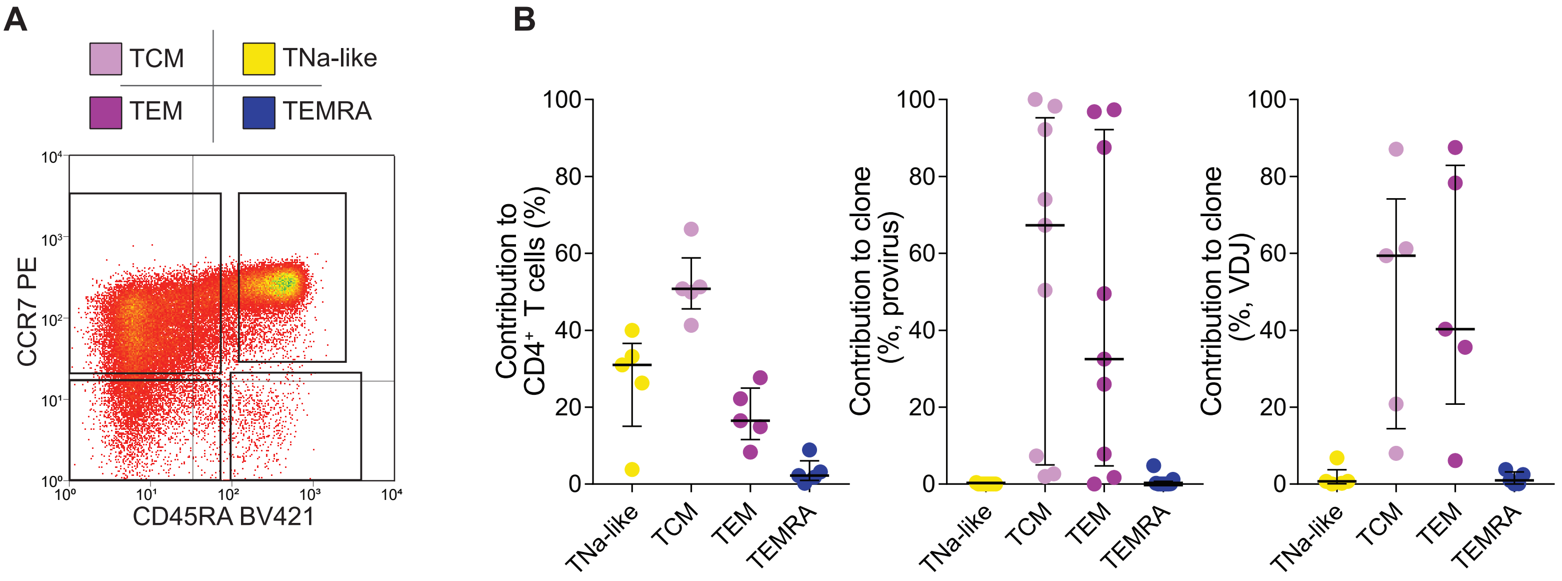


**Supplementary Figure S10. Contribution of $CD4^+$ T cell subsets to antigen-responsive clonotypes carrying HIV proviruses.**

(**A**) Representative experiment and sorting strategy to isolate $CD4^+$ T cell subsets previously gated on single, live, $CD14^-CD16^-CD20^-$, $CD3^+CD4^+$ cells. (**B**) Relative contribution (%) of $CD4^+$ T cell subsets to total $CD4^+$ T cells (left panel) and to proviral copies (center panel) and VDJ rearrangement copies (right panel) of antigen-responsive HIV-1-infected clones.

**Supplementary Table S1.** Participant characteristics.

| Participant ID | Sex | Race | Age | HIV-RNA pre ART ($log_{10}$ cps/mL) | HIV-RNA current ($log_{10}$ cps/mL) | $CD4^+$ T cells nadir (cells/$mm^3$) | $CD4^+$ T cells current (cells/$mm^3$) | $CD4^+$ T cell recovery (cells/$mm^3$) | Years since HIV-1 diagnosis | Years on ART[a] | Years with <50 HIV-RNA cps/mL[b] | Current ART Regimen[c] |
|---|---|---|---|---|---|---|---|---|---|---|---|---|
| P1 | Male | White | 56 | 5.1 | <1.3 | 328 | 1429 | 1101 | 19.8 | 5.9 | 4.8 | ABC/ 3TC/DTG |
| P2 | Male | AA | 57 | 5.0 | <1.3 | 158 | 630 | 472 | 24.4 | 24.4 | 11.9 | ZDV/3TC/EFV/ATZ/r |
| P3 | Female | AA | 53 | 4.1 | <1.3 | 320 | 1224 | 904 | 19.5 | 19.5 | 10.2 | ABC/3TC/DTG |
| P4 | Female | AA | 60 | 4.7 | <1.3 | 159 | 475 | 316 | 13.3 | 11.0 | 8.0 | TAF/FTC/ATV/c |
| P5 | Male | AA | 54 | 4.7 | <1.3 | 184 | 1194 | 1010 | 21.0 | 12.9 | 6.5 | TDF/FTC/RPV |
| P6 | Female | AA | 39 | 4.7 | <1.3 | 275 | 1098 | 823 | 25.4 | 6.4 | 6.1 | ABC/3TC/DTG |
| P7 | Female | AA | 59 | 4.9 | <1.3 | 81 | 654 | 573 | 24.4 | 8.8 | 4.4 | ETV/RAL/DRV/r |
| P8 | Male | White | 63 | 5.5 | <1.3 | 41 | 407 | 366 | 16.9 | 16.5 | 16.2 | TAF/FTC/RPV |
| P9 | Male | White | 65 | 4.5 | <1.6 | 105 | 518 | 413 | 28.7 | 28.0 | 11.8 | TAF/FTC/RPV/DTG |
| P10 | Male | White | 59 | 5.9 | <1.6 | 205 | 516 | 311 | 28.4 | 27.8 | 7.9 | ABC/3TC/DTG |
| Median | | | 58 | 4.8 | <1.3 | 171.5 | 642 | 522.5 | 22.7 | 14.7 | 7.9 | |
| IQR | | | 53.7-60. | 4.6-5.2 | | 99-286 | 505.8-1202 | 353.5-930.5 | 18.8-26.1 | 8.2-25.5 | 5.7-11.8 | |

[a]Years since the beginning of first ART regimen

[b]Years of stable suppression with plasma HIV-RNA below limit of detection of clinical assays

[c]Antiretroviral drug abbreviations: ABC (abacavir); ATV/r (atazanavir/ritonavir); ATV/c (aazanavir/cobicistat); DRV/r (darunavir/ritonavir); DTG (dolutegravir); EFV (efavirenz); ETV (etravirine); FTC (emtricitabine); TDF (tenofovir disoproxil fumarate); RAL (raltegravir); ZDV (zidovudine); 3TC (lamivudine)

**Supplementary Table S2.** Single genome sequences and oligoclonality of proviral populations from sorted CD4+ T cells.

| | | CMV-stim CD40L+CD69+CD4+ cells | | | Gag-stim CD40L+CD69+CD4+ cells | | | CD3/28-stim CD40L+CD69+CD4+ cells | | | CMV non responding $CD45RO^{hi}CD4+$ cells | | | Gag non responding $CD45RO^{hi}CD4+$ cells | | |
|---|---|---|---|---|---|---|---|---|---|---|---|---|---|---|---|---|
| Participant ID | Amplicon | Total | Identical (%) | Gini | Total | Identical (%) | Gini | Total | Identical (%) | Gini | Total | Identical (%) | Gini | Total | Identical (%) | Gini |
| P1 | *u5-gag* | 66 | 48 (73) | 0.64 | 34 | 12 (35) | 0.25 | 42 | 8 (19) | 0.09 | | | | | | |
| P2 | *u5-gag* | 34 | 22 (65) | 0.48 | 24 | 16 (67) | 0.41 | 61 | 13 (21) | 0.16 | 39 | 26 (67) | 0.55 | 28 | 20 (71) | 0.51 |
| P3 | *u5-gag* | 88 | 60 (68) | 0.50 | 16 | 7 (44) | 0.28 | 73 | 25 (34) | 0.18 | 41 | 6 (15) | 0.07 | 22 | 0 (0) | 0.00 |
| P4 | *u5-gag* | 57 | 51 (89) | 0.77 | 40 | 6 (15) | 0.10 | 46 | 18 (39) | 0.27 | | | | | | |
| P5 | *env* | 33 | 28 (85) | 0.70 | 33 | 10 (30) | 0.13 | 50 | 37 (74) | 0.64 | 75 | 36 (48) | 0.38 | 55 | 42 (76) | 0.68 |
| P5 | *u5-gag* | 11 | 9 (82) | 0.43 | | | | 22 | 9 (41) | 0.25 | 9 | 2 (22) | 0.11 | 8 | 3 (38) | 0.24 |
| P6 | *u5-gag* | 12 | 3 (25) | 0.16 | 55 | 15 (27) | 0.17 | 19 | 0 (0) | 0.00 | 52 | 9 (17) | 0.09 | 33 | 2 (6) | 0.03 |
| P7 | *u5-gag* | 22 | 14 (64) | 0.37 | 67 | 35 (52) | 0.35 | 41 | 6 (15) | 0.07 | | | | 52* | 28 (54)* | 0.32* |
| P8 | *u5-gag* | 123 | 111 (90) | 0.83 | | | | 59 | 24 (41) | 0.29 | 72 | 14 (19) | 0.12 | | | |
| P9 | *env* | 45 | 35 (78) | 0.51 | | | | 15 | 4 (27) | 0.12 | 49 | 26 (53) | 0.36 | | | |
| P10 | *u5-gag* | 14 | 3 (21) | 0.14 | 24 | 7 (29) | 0.20 | 26 | 0 (0) | 0.00 | | | | | | |
| Total | | 505 | 384 (76) | | 293 | 108 (37) | | 454 | 144 (32) | | 337 | 119 (35) | | 146 | 67 (46) | |
| Average | | 46 | 35 (67) | 0.50 | 37 | 14 (37) | 0.24 | 41 | 13 (28) | 0.19 | 48 | 17 (34) | 0.24 | 29 | 13 (24) | 0.29 |

* CMV and Gag non responding $CD45RO^{hi}$ $CD4^{+}$ cells were pooled, and excluded from calculations

**Supplementary Table S3.** Integration site analysis of HIV-1-infected antigen-responding clones.

| **Sample** | | | **Provirus** | | | | | | | | **Host gene** | | | | |
|---|---|---|---|---|---|---|---|---|---|---|---|---|---|---|---|
| **Participant ID** | **Antigen** | **Time point (months)[a]** | **Provirus ID** | **Chr.** | **Position[b]** | **5nt duplication[c]** | **Host junction recovered** | **Strand** | **Orientation relative to gene** | **Method[d]** | **Gene symbol[e]** | **Gene ID** | **Gene transcription orientation** | **Intron/exon/ distance[f]** | **Identified before (ref)[g]** |
| P1 | CMV | 0,4,6,7 | 1.c.mkl1 | 22 | 40557594 | GTCTT | 5', 3' | - | same | B, C | ***MKL1*** ~ | 57591 | - | intron 2 | 8, 9, 10, 12 |
| | CMV | 0,4,6 | 1.c.memo1 | 2 | 31948508 | ATTTC | 5', 3' | - | same | B, C | *MEMO1* | 51072 | - | intron 1 | 12 |
| P2 | CMV | 0,2,11 | 2.c.stat5b | 17 | 42241492 | CTCCA | 5', 3' | - | same | A, C | ***STAT5B*** ~ | 6777 | - | intron 1 | 8, 9, 10, 47 |
| | CMV | 0,2 | 2.c.ddi2 | 1 | 15658310 | GGACC | 5' | - | opposite | A | *DDI2* | 84301 | + | intron 9 | 12 |
| | CMV | 0,2,11 | 2.c.hivep3 | 1 | 41985926 | AATAT | 5' | + | opposite | A, C | *HIVEP3*~ | 59269 | - | intron 1 | 8, 10, 12 |
| | CMV | 0,2 | 2.c.mir194 | 11 | 64885345 | CCAAT | 3' | + | opposite | A | *MIR194-2HG* | 105369343 | - | ig, 3112 | |
| P3 | CMV | 0,3,7,8,10 | 3.c.fbxo22 | 15 | 75914981 | AAAAC | 5', 3' | - | opposite | A, B, C | *FBXO22* | 26263 | + | intron 4 | 12 |
| | CMV | 0,7 | 3.c.gdap2 | 1 | 117864268 | CTTCC | 5', 3' | + | opposite | A, C | *GDAP2* | 54834 | - | exon 14 | 12 |
| | CMV | 0,3,7,8,10 | 3.c.ccnl2 | 1 | 1390336 | CTTTT | 5' | + | opposite | A | *CCNL2* | 81669 | - | intron 8 | 10, 12 |
| | CMV | 0,3,7,8,10 | 3.c.bach2 | 6 | 90052250 | GCTTG | 5', 3' | - | same | A, B, C | ***BACH2*** ~ | 60468 | - | intron 5 | 8, 9, 10, 12, 47 |
| P4 | CMV | 0,5,6 | 4.c.st6 | 1 | 76337191 | CCCGC/ab | 5', 3' | + | same | B, C | *ST6GALNAC3* | 256435 | + | intron 2 | 8, 12 |
| P5 | CMV | 0,4,10 | 5.c.kcnc2 | 12 | 74917002 | GTTTT | 5', 3' | + | opposite | A, B, C | *KCNC2* | 3747 | - | ig, 123075 | |
| | CMV | 0,4,10 | 5.c.dec1 | 9 | 114949387 | GGTGT | 5', 3' | - | opposite | A, B, C | *DELEC1*~ | 50514 | + | intron 2 | |
| P6 | CMV | 0,19 | 6.c.nol2 | 2 | 10680857 | CACTC | 3' | + | opposite | B, C | *NOL10* | 79954 | - | intron 3 | 8 |
| | Gag | 0,12,25 | 6.g.bach2.c1 | 6 | 90230775 | GTTTC | 5', 3' | - | same | A, B, C | ***BACH2*** ~ | 60468 | - | intron 3 | 8, 9, 10, 12, 47 |
| | Gag | 0,12,25 | 6.g.bach2.c2 | 6 | 90063562 | GGTAT/ab | 5', 3' | - | same | A, B, C | ***BACH2*** ~ | 60468 | - | intron 5 | 8, 9, 10, 12, 47 |
| | Gag | 0,12,25 | 6.g.bach2.c3 | 6 | 90037792 | CCCAT | 5', 3' | - | same | A, B, C | ***BACH2*** ~ | 60468 | - | intron 5 | 8, 9, 10, 12, 47 |
| | Gag | 0,12,25 | 6.g.kpna2 | 1 | 32131583 | TATAT | 5', 3' | - | opposite | A, B, C | *KPNA6* | 3838 | + | intron 1 | 10 |
| | Gag | 0,12,25 | 6.g.cramp1 | 16 | 1650160 | AATAC | 5', 3' | - | opposite | A, C | *CRAMP1* | 57585 | + | intron 6 | 9, 12 |
| | Gag | 0,12,25 | 6.g.stat5b | 17 | 42234035 | ATAAC | 5', 3' | - | same | A, C | ***STAT5B*** ~ | 6777 | - | intron 1 | 9, 47 |
| P8 | CMV | 0,5 | 8.c.paf | 17 | 2671849 | GCGGG | 5' | - | opposite | A | *PAFAH1B1* | 5048 | + | intron 6 | 8, 12 |
| | CMV | 0,5 | 8.c.dle | 13 | 50070306 | TAAAC | 5', 3' | + | opposite | A | *DLEU2*~ | 8847 | - | intron 4 | 12 |

a Provirus detected by sequence identity or integration site
b Number refers to the third nucleotide of the duplication, using GRCH38/hg38 assembly
c Insertion junction not at the beginning or end of the LTR is defined as aberrant (ab)
d A (INSPIIRED, LM-PCR plus paired-end sequencing), B (Lenti-X, LM-PCR plus direct Sanger), C (ISS-PCR, standard or dd PCR using primers across the integration junction)
e Genes previosuly linked to HIV-1 persistence are highlighted in red; ~ indicates the gene is a cancer related gene
f If integration site is intergenic (ig) nucleotide distance from nearest gene is provided
g Data from NCI Retrovirus Integration Database[87] ; only studies on ART-treated individuals were included

**Supplementary Table S4.** Characteristics of TCRβ sequences in clusters shown in Figure 4.

Characteristics of the TCR sequences within clusters displayed in Figure 4. HLA genes with alleles shared among all subjects within a cluster are highlighted in pink. Predicted binding HLA alleles, significantly enriched in subjects within the cluster, are highlighted in yellow.

| TCR Cluster | Participant | Stimulation | TCRβ | V chain | J chain | Templates | HLA-DPA1 | HLA-DPB1 | HLA-DQA1 | HLA-DQB1 | HLA-DRB1 | HLA-DRB3 | HLA-DRB4 | HLA-DRB5 |
|---|---|---|---|---|---|---|---|---|---|---|---|---|---|---|
| Group 1 - CMV - %GSTE - global | P1 | CMV lysate | CASVGSTEAFF | TRBV19-01 | TRBJ01-01 | 224 | *01:03/*02:01 | *104:01/*11:01 | *02:01/*05:05 | *02:02/*03:01 | *07:01/*13:03 | *01:01 | *01:01 | - |
| | P1 | CMV lysate | CASVGSTEAFF | TRBV27-01 | TRBJ01-01 | 5 | *01:03/*02:01 | *104:01/*11:01 | *02:01/*05:05 | *02:02/*03:01 | *07:01/*13:03 | *01:01 | *01:01 | - |
| | P1 | CMV lysate | CATTGSTEAFF | TRBV19-01 | TRBJ01-01 | 1 | *01:03/*02:01 | *104:01/*11:01 | *02:01/*05:05 | *02:02/*03:01 | *07:01/*13:03 | *01:01 | *01:01 | - |
| | P1 | CMV lysate | CASGGSTEAFF | TRBV19-01 | TRBJ01-01 | 20 | *01:03/*02:01 | *104:01/*11:01 | *02:01/*05:05 | *02:02/*03:01 | *07:01/*13:03 | *01:01 | *01:01 | - |
| | P1 | CMV lysate | CASGGSTEAFF | TRBV19-01 | TRBJ01-01 | 17 | *01:03/*02:01 | *104:01/*11:01 | *02:01/*05:05 | *02:02/*03:01 | *07:01/*13:03 | *01:01 | *01:01 | - |
| | P1 | CMV lysate | CASGGSTEAFF | TRBV19-01 | TRBJ01-01 | 2 | *01:03/*02:01 | *104:01/*11:01 | *02:01/*05:05 | *02:02/*03:01 | *07:01/*13:03 | *01:01 | *01:01 | - |
| | P1 | CMV lysate | CASGGSTEAFF | TRBV19-01 | TRBJ01-01 | 1 | *01:03/*02:01 | *104:01/*11:01 | *02:01/*05:05 | *02:02/*03:01 | *07:01/*13:03 | *01:01 | *01:01 | - |
| | P9 | CMV lysate | CASGGSTEAFF | TRBV19-01 | TRBJ01-01 | 11 | - | *04:01/*04:01 | *01:01/*06:01 | *03:01/*05:01 | *01:03/*08:03 | - | - | - |
| | P1 | CMV lysate | CASSGSTEAFF | TRBV27-01 | TRBJ01-01 | 360 | *01:03/*02:01 | *104:01/*11:01 | *02:01/*05:05 | *02:02/*03:01 | *07:01/*13:03 | *01:01 | *01:01 | - |
| | P1 | CMV lysate | CASSGSTEAFF | TRBV27-01 | TRBJ01-01 | 51 | *01:03/*02:01 | *104:01/*11:01 | *02:01/*05:05 | *02:02/*03:01 | *07:01/*13:03 | *01:01 | *01:01 | - |
| | P1 | CMV lysate | CASSGSTEAFF | TRBV27-01 | TRBJ01-01 | 17 | *01:03/*02:01 | *104:01/*11:01 | *02:01/*05:05 | *02:02/*03:01 | *07:01/*13:03 | *01:01 | *01:01 | - |
| | P1 | CMV lysate | CASSGSTEAFF | TRBV19-01 | TRBJ01-01 | 3 | *01:03/*02:01 | *104:01/*11:01 | *02:01/*05:05 | *02:02/*03:01 | *07:01/*13:03 | *01:01 | *01:01 | - |
| | P1 | CMV lysate | CASSGSTEAFF | TRBV27-01 | TRBJ01-01 | 2 | *01:03/*02:01 | *104:01/*11:01 | *02:01/*05:05 | *02:02/*03:01 | *07:01/*13:03 | *01:01 | *01:01 | - |
| | P1 | CMV lysate | CASSGSTEAFF | TRBV27-01 | TRBJ01-01 | 1 | *01:03/*02:01 | *104:01/*11:01 | *02:01/*05:05 | *02:02/*03:01 | *07:01/*13:03 | *01:01 | *01:01 | - |
| | P9 | CMV lysate | CASSGSTEAFF | TRBV19-01 | TRBJ01-01 | 2 | - | *04:01/*04:01 | *01:01/*06:01 | *03:01/*05:01 | *01:03/*08:03 | - | - | - |
| | P9 | CMV lysate | CASSGSTEQYF | TRBV19-01 | TRBJ02-07 | 1 | - | *04:01/*04:01 | *01:01/*06:01 | *03:01/*05:01 | *01:03/*08:03 | - | - | - |
| | P1 | CMV lysate | CATSGSTEAFF | TRBV19-01 | TRBJ01-01 | 1 | *01:03/*02:01 | *104:01/*11:01 | *02:01/*05:05 | *02:02/*03:01 | *07:01/*13:03 | *01:01 | *01:01 | - |
| | P9 | CMV lysate | CATSGSTEAFF | TRBV03-01/03-02 | TRBJ01-01 | 1 | - | *04:01/*04:01 | *01:01/*06:01 | *03:01/*05:01 | *01:03/*08:03 | - | - | - |
| | P1 | CMV lysate | CGSGGSTEAFF | TRBV19-01 | TRBJ01-01 | 1 | *01:03/*02:01 | *104:01/*11:01 | *02:01/*05:05 | *02:02/*03:01 | *07:01/*13:03 | *01:01 | *01:01 | - |
| | P1 | CMV lysate | CASTGSTEAFF | TRBV27-01 | TRBJ01-01 | 170 | *01:03/*02:01 | *104:01/*11:01 | *02:01/*05:05 | *02:02/*03:01 | *07:01/*13:03 | *01:01 | *01:01 | - |
| | P1 | CMV lysate | CASTGSTEAFF | TRBV19-01 | TRBJ01-01 | 123 | *01:03/*02:01 | *104:01/*11:01 | *02:01/*05:05 | *02:02/*03:01 | *07:01/*13:03 | *01:01 | *01:01 | - |
| | P1 | CMV lysate | CASTGSTEAFF | TRBV19-01 | TRBJ01-01 | 28 | *01:03/*02:01 | *104:01/*11:01 | *02:01/*05:05 | *02:02/*03:01 | *07:01/*13:03 | *01:01 | *01:01 | - |
| | P1 | CMV lysate | CASTGSTEAFF | TRBV27-01 | TRBJ01-01 | 19 | *01:03/*02:01 | *104:01/*11:01 | *02:01/*05:05 | *02:02/*03:01 | *07:01/*13:03 | *01:01 | *01:01 | - |
| | P1 | CMV lysate | CASTGSTEAFF | TRBV27-01 | TRBJ01-01 | 8 | *01:03/*02:01 | *104:01/*11:01 | *02:01/*05:05 | *02:02/*03:01 | *07:01/*13:03 | *01:01 | *01:01 | - |
| | P1 | CMV lysate | CASTGSTEAFF | TRBV03-01/03-02 | TRBJ01-01 | 1 | *01:03/*02:01 | *104:01/*11:01 | *02:01/*05:05 | *02:02/*03:01 | *07:01/*13:03 | *01:01 | *01:01 | - |
| | P1 | CMV lysate | CASTGSTEAFF | TRBV19-01 | TRBJ01-01 | 1 | *01:03/*02:01 | *104:01/*11:01 | *02:01/*05:05 | *02:02/*03:01 | *07:01/*13:03 | *01:01 | *01:01 | - |
| | P4 | CMV lysate | CASTGSTEAFF | TRBV19-01 | TRBJ01-01 | 21 | *01:03/*04:01 | - | *01:02/*06:01 | *03:01/*06:02 | *08:03/*15:01 | - | - | *01:01 |
| | P4 | CMV lysate | CASTGSTEAFF | TRBV19-01 | TRBJ01-01 | 14 | *01:03/*04:01 | - | *01:02/*06:01 | *03:01/*06:02 | *08:03/*15:01 | - | - | *01:01 |
| | P4 | CMV lysate | CASTGSTEAFF | TRBV19-01 | TRBJ01-01 | 13 | *01:03/*04:01 | - | *01:02/*06:01 | *03:01/*06:02 | *08:03/*15:01 | - | - | *01:01 |
| | P4 | CMV lysate | CASTGSTEAFF | TRBV19-01 | TRBJ01-01 | 4 | *01:03/*04:01 | - | *01:02/*06:01 | *03:01/*06:02 | *08:03/*15:01 | - | - | *01:01 |
| | P9 | CMV lysate | CASTGSTEAFF | TRBV19-01 | TRBJ01-01 | 40 | - | *04:01/*04:01 | *01:01/*06:01 | *03:01/*05:01 | *01:03/*08:03 | - | - | - |
| | P9 | CMV lysate | CASTGSTEAFF | TRBV19-01 | TRBJ01-01 | 2 | - | *04:01/*04:01 | *01:01/*06:01 | *03:01/*05:01 | *01:03/*08:03 | - | - | - |
| | P9 | CMV lysate | CASTGSTEAFF | TRBV27-01 | TRBJ01-01 | 1 | - | *04:01/*04:01 | *01:01/*06:01 | *03:01/*05:01 | *01:03/*08:03 | - | - | - |
| | P9 | CMV lysate | CASTGSTEAFF | TRBV19-01 | TRBJ01-01 | 1 | - | *04:01/*04:01 | *01:01/*06:01 | *03:01/*05:01 | *01:03/*08:03 | - | - | - |
| | P1 | CMV lysate | CASCGSTEAFF | TRBV19-01 | TRBJ01-01 | 1 | *01:03/*02:01 | *104:01/*11:01 | *02:01/*05:05 | *02:02/*03:01 | *07:01/*13:03 | *01:01 | *01:01 | - |
| | P4 | CMV lysate | CASFGSTEAFF | TRBV27-01 | TRBJ01-01 | 1 | *01:03/*04:01 | - | *01:02/*06:01 | *03:01/*06:02 | *08:03/*15:01 | - | - | *01:01 |
| | P1 | CMV lysate | CASAGSTEAFF | TRBV12-05 | TRBJ01-01 | 164 | *01:03/*02:01 | *104:01/*11:01 | *02:01/*05:05 | *02:02/*03:01 | *07:01/*13:03 | *01:01 | *01:01 | - |
| | P1 | CMV lysate | CASAGSTEAFF | TRBV19-01 | TRBJ01-01 | 18 | *01:03/*02:01 | *104:01/*11:01 | *02:01/*05:05 | *02:02/*03:01 | *07:01/*13:03 | *01:01 | *01:01 | - |
| | P4 | CMV lysate | CASAGSTEAFF | TRBV19-01 | TRBJ01-01 | 6 | *01:03/*04:01 | - | *01:02/*06:01 | *03:01/*06:02 | *08:03/*15:01 | - | - | *01:01 |
| | P1 | CMV lysate | CAAAGSTEAFF | TRBV27-01 | TRBJ01-01 | 4 | *01:03/*02:01 | *104:01/*11:01 | *02:01/*05:05 | *02:02/*03:01 | *07:01/*13:03 | *01:01 | *01:01 | - |
| | P1 | CMV lysate | CASLGSTEAFF | TRBV27-01 | TRBJ01-01 | 187 | *01:03/*02:01 | *104:01/*11:01 | *02:01/*05:05 | *02:02/*03:01 | *07:01/*13:03 | *01:01 | *01:01 | - |
| | P1 | CMV lysate | CASLGSTEAFF | TRBV06-05 | TRBJ01-01 | 2 | *01:03/*02:01 | *104:01/*11:01 | *02:01/*05:05 | *02:02/*03:01 | *07:01/*13:03 | *01:01 | *01:01 | - |
| | P1 | CMV lysate | CASIGSTEAFF | TRBV27-01 | TRBJ01-01 | 309 | *01:03/*02:01 | *104:01/*11:01 | *02:01/*05:05 | *02:02/*03:01 | *07:01/*13:03 | *01:01 | *01:01 | - |
| | P1 | CMV lysate | CASIGSTEAFF | TRBV19-01 | TRBJ01-01 | 2 | *01:03/*02:01 | *104:01/*11:01 | *02:01/*05:05 | *02:02/*03:01 | *07:01/*13:03 | *01:01 | *01:01 | - |
| | P1 | CMV lysate | CASIGSTEAFF | TRBV06-05 | TRBJ01-01 | 2 | *01:03/*02:01 | *104:01/*11:01 | *02:01/*05:05 | *02:02/*03:01 | *07:01/*13:03 | *01:01 | *01:01 | - |
| | P4 | CMV lysate | CASIGSTEAFF | TRBV19-01 | TRBJ01-01 | 2 | *01:03/*04:01 | - | *01:02/*06:01 | *03:01/*06:02 | *08:03/*15:01 | - | - | *01:01 |
| | P9 | CMV lysate | CASIGSTEAFF | TRBV02-01 | TRBJ01-01 | 8 | - | *04:01/*04:01 | *01:01/*06:01 | *03:01/*05:01 | *01:03/*08:03 | - | - | - |
| | P1 | CMV lysate | CASMGSTEAFF | TRBV19-01 | TRBJ01-01 | 2 | *01:03/*02:01 | *104:01/*11:01 | *02:01/*05:05 | *02:02/*03:01 | *07:01/*13:03 | *01:01 | *01:01 | - |
| | P4 | CMV lysate | CASMGSTEAFF | TRBV19-01 | TRBJ01-01 | 1 | *01:03/*04:01 | - | *01:02/*06:01 | *03:01/*06:02 | *08:03/*15:01 | - | - | *01:01 |
| | P9 | CMV lysate | CASMGSTEAFF | TRBV19-01 | TRBJ01-01 | 49 | - | *04:01/*04:01 | *01:01/*06:01 | *03:01/*05:01 | *01:03/*08:03 | - | - | - |
| | P9 | CMV lysate | CASMGSTEAFF | TRBV19-01 | TRBJ01-01 | 11 | - | *04:01/*04:01 | *01:01/*06:01 | *03:01/*05:01 | *01:03/*08:03 | - | - | - |
| | P1 | CMV lysate | CACRGSTEAFF | TRBV03-01/03-02 | TRBJ01-01 | 36 | *01:03/*02:01 | *104:01/*11:01 | *02:01/*05:05 | *02:02/*03:01 | *07:01/*13:03 | *01:01 | *01:01 | - |
| | P9 | CMV lysate | CASRGSTEQYF | TRBV27-01 | TRBJ02-07 | 1 | - | *04:01/*04:01 | *01:01/*06:01 | *03:01/*05:01 | *01:03/*08:03 | - | - | - |
| | P1 | CMV lysate | CASRGSTEAFF | TRBV03-01/03-02 | TRBJ01-01 | 3111 | *01:03/*02:01 | *104:01/*11:01 | *02:01/*05:05 | *02:02/*03:01 | *07:01/*13:03 | *01:01 | *01:01 | - |
| | P1 | CMV lysate | CASRGSTEAFF | TRBV19-01 | TRBJ01-01 | 1747 | *01:03/*02:01 | *104:01/*11:01 | *02:01/*05:05 | *02:02/*03:01 | *07:01/*13:03 | *01:01 | *01:01 | - |
| | P1 | CMV lysate | CASRGSTEAFF | TRBV27-01 | TRBJ01-01 | 1061 | *01:03/*02:01 | *104:01/*11:01 | *02:01/*05:05 | *02:02/*03:01 | *07:01/*13:03 | *01:01 | *01:01 | - |
| | P1 | CMV lysate | CASRGSTEAFF | TRBV19-01 | TRBJ01-01 | 509 | *01:03/*02:01 | *104:01/*11:01 | *02:01/*05:05 | *02:02/*03:01 | *07:01/*13:03 | *01:01 | *01:01 | - |
| | P1 | CMV lysate | CASRGSTEAFF | TRBV03-01/03-02 | TRBJ01-01 | 184 | *01:03/*02:01 | *104:01/*11:01 | *02:01/*05:05 | *02:02/*03:01 | *07:01/*13:03 | *01:01 | *01:01 | - |
| | P1 | CMV lysate | CASRGSTEAFF | TRBV03-01/03-02 | TRBJ01-01 | 100 | *01:03/*02:01 | *104:01/*11:01 | *02:01/*05:05 | *02:02/*03:01 | *07:01/*13:03 | *01:01 | *01:01 | - |
| | P1 | CMV lysate | CASRGSTEAFF | TRBV27-01 | TRBJ01-01 | 8 | *01:03/*02:01 | *104:01/*11:01 | *02:01/*05:05 | *02:02/*03:01 | *07:01/*13:03 | *01:01 | *01:01 | - |
| | P1 | CMV lysate | CASRGSTEAFF | TRBV03-01/03-02 | TRBJ01-01 | 6 | *01:03/*02:01 | *104:01/*11:01 | *02:01/*05:05 | *02:02/*03:01 | *07:01/*13:03 | *01:01 | *01:01 | - |
| | P1 | CMV lysate | CASRGSTEAFF | TRBV27-01 | TRBJ01-01 | 4 | *01:03/*02:01 | *104:01/*11:01 | *02:01/*05:05 | *02:02/*03:01 | *07:01/*13:03 | *01:01 | *01:01 | - |
| | P1 | CMV lysate | CASRGSTEAFF | TRBV19-01 | TRBJ01-01 | 3 | *01:03/*02:01 | *104:01/*11:01 | *02:01/*05:05 | *02:02/*03:01 | *07:01/*13:03 | *01:01 | *01:01 | - |
| | P1 | CMV lysate | CASRGSTEAFF | TRBV03-01/03-02 | TRBJ01-01 | 1 | *01:03/*02:01 | *104:01/*11:01 | *02:01/*05:05 | *02:02/*03:01 | *07:01/*13:03 | *01:01 | *01:01 | - |
| | P1 | CMV lysate | CASRGSTEAFF | TRBV27-01 | TRBJ01-01 | 1 | *01:03/*02:01 | *104:01/*11:01 | *02:01/*05:05 | *02:02/*03:01 | *07:01/*13:03 | *01:01 | *01:01 | - |
| | P1 | CMV lysate | CASRGSTEAFF | TRBV19-01 | TRBJ01-01 | 1 | *01:03/*02:01 | *104:01/*11:01 | *02:01/*05:05 | *02:02/*03:01 | *07:01/*13:03 | *01:01 | *01:01 | - |
| | P1 | CMV lysate | CASRGSTEAFF | TRBV19-01 | TRBJ01-01 | 1 | *01:03/*02:01 | *104:01/*11:01 | *02:01/*05:05 | *02:02/*03:01 | *07:01/*13:03 | *01:01 | *01:01 | - |
| | P1 | CMV lysate | CASRGSTEAFF | TRBV27-01 | TRBJ01-01 | 1 | *01:03/*02:01 | *104:01/*11:01 | *02:01/*05:05 | *02:02/*03:01 | *07:01/*13:03 | *01:01 | *01:01 | - |
| | P1 | CMV lysate | CASRGSTEAFF | TRBV19-01 | TRBJ01-01 | 1 | *01:03/*02:01 | *104:01/*11:01 | *02:01/*05:05 | *02:02/*03:01 | *07:01/*13:03 | *01:01 | *01:01 | - |
| | P1 | CMV lysate | CASRGSTEAFF | TRBV19-01 | TRBJ01-01 | 1 | *01:03/*02:01 | *104:01/*11:01 | *02:01/*05:05 | *02:02/*03:01 | *07:01/*13:03 | *01:01 | *01:01 | - |
| | P4 | CMV lysate | CASRGSTEAFF | TRBV19-01 | TRBJ01-01 | 1 | *01:03/*04:01 | - | *01:02/*06:01 | *03:01/*06:02 | *08:03/*15:01 | - | - | *01:01 |
| | P4 | CMV lysate | CASRGSTEAFF | TRBV19-01 | TRBJ01-01 | 1 | *01:03/*04:01 | - | *01:02/*06:01 | *03:01/*06:02 | *08:03/*15:01 | - | - | *01:01 |
| | P4 | CMV lysate | CASRGSTEAFF | TRBV27-01 | TRBJ01-01 | 1 | *01:03/*04:01 | - | *01:02/*06:01 | *03:01/*06:02 | *08:03/*15:01 | - | - | *01:01 |
| | P9 | CMV lysate | CASRGSTEAFF | TRBV27-01 | TRBJ01-01 | 28 | - | *04:01/*04:01 | *01:01/*06:01 | *03:01/*05:01 | *01:03/*08:03 | - | - | - |
| | P9 | CMV lysate | CASRGSTEAFF | TRBV27-01 | TRBJ01-01 | 5 | - | *04:01/*04:01 | *01:01/*06:01 | *03:01/*05:01 | *01:03/*08:03 | - | - | - |
| | P9 | CMV lysate | CASRGSTEAFF | TRBV19-01 | TRBJ01-01 | 3 | - | *04:01/*04:01 | *01:01/*06:01 | *03:01/*05:01 | *01:03/*08:03 | - | - | - |
| | P4 | CMV lysate | CVTRGSTEAFF | TRBV19-01 | TRBJ01-01 | 1 | *01:03/*04:01 | - | *01:02/*06:01 | *03:01/*06:02 | *08:03/*15:01 | - | - | *01:01 |
| | P1 | CMV lysate | CASYGSTEAFF | TRBV19-01 | TRBJ01-01 | 1 | *01:03/*02:01 | *104:01/*11:01 | *02:01/*05:05 | *02:02/*03:01 | *07:01/*13:03 | *01:01 | *01:01 | - |
| | P1 | CMV lysate | CATRGSTEAFF | TRBV27-01 | TRBJ01-01 | 592 | *01:03/*02:01 | *104:01/*11:01 | *02:01/*05:05 | *02:02/*03:01 | *07:01/*13:03 | *01:01 | *01:01 | - |
| | P1 | CMV lysate | CATRGSTEAFF | TRBV03-01/03-02 | TRBJ01-01 | 90 | *01:03/*02:01 | *104:01/*11:01 | *02:01/*05:05 | *02:02/*03:01 | *07:01/*13:03 | *01:01 | *01:01 | - |
| | P1 | CMV lysate | CATRGSTEAFF | TRBV03-01/03-02 | TRBJ01-01 | 68 | *01:03/*02:01 | *104:01/*11:01 | *02:01/*05:05 | *02:02/*03:01 | *07:01/*13:03 | *01:01 | *01:01 | - |
| | P1 | CMV lysate | CATRGSTEAFF | TRBV27-01 | TRBJ01-01 | 24 | *01:03/*02:01 | *104:01/*11:01 | *02:01/*05:05 | *02:02/*03:01 | *07:01/*13:03 | *01:01 | *01:01 | - |
| | P1 | CMV lysate | CATRGSTEAFF | TRBV27-01 | TRBJ01-01 | 9 | *01:03/*02:01 | *104:01/*11:01 | *02:01/*05:05 | *02:02/*03:01 | *07:01/*13:03 | *01:01 | *01:01 | - |
| | P1 | CMV lysate | CATRGSTEAFF | TRBV19-01 | TRBJ01-01 | 2 | *01:03/*02:01 | *104:01/*11:01 | *02:01/*05:05 | *02:02/*03:01 | *07:01/*13:03 | *01:01 | *01:01 | - |
| | P1 | CMV lysate | CASQGSTEAFF | TRBV19-01 | TRBJ01-01 | 83 | *01:03/*02:01 | *104:01/*11:01 | *02:01/*05:05 | *02:02/*03:01 | *07:01/*13:03 | *01:01 | *01:01 | - |
| | P1 | CMV lysate | CASQGSTEAFF | TRBV27-01 | TRBJ01-01 | 3 | *01:03/*02:01 | *104:01/*11:01 | *02:01/*05:05 | *02:02/*03:01 | *07:01/*13:03 | *01:01 | *01:01 | - |
| | P9 | CMV lysate | CASQGSTEAFF | TRBV19-01 | TRBJ01-01 | 1 | - | *04:01/*04:01 | *01:01/*06:01 | *03:01/*05:01 | *01:03/*08:03 | - | - | - |
| | P9 | CMV lysate | CASQGSTEQFF | TRBV19-01 | TRBJ02-01 | 12 | - | *04:01/*04:01 | *01:01/*06:01 | *03:01/*05:01 | *01:03/*08:03 | - | - | - |
| | P4 | CMV lysate | CATQGSTEAFF | TRBV03-01/03-02 | TRBJ01-01 | 1 | *01:03/*04:01 | - | *01:02/*06:01 | *03:01/*06:02 | *08:03/*15:01 | - | - | *01:01 |
| | P1 | CMV lysate | CASKGSTEAFF | TRBV19-01 | TRBJ01-01 | 254 | *01:03/*02:01 | *104:01/*11:01 | *02:01/*05:05 | *02:02/*03:01 | *07:01/*13:03 | *01:01 | *01:01 | - |
| | P1 | CMV lysate | CASKGSTEAFF | TRBV19-01 | TRBJ01-01 | 80 | *01:03/*02:01 | *104:01/*11:01 | *02:01/*05:05 | *02:02/*03:01 | *07:01/*13:03 | *01:01 | *01:01 | - |
| | P1 | CMV lysate | CASKGSTEAFF | TRBV19-01 | TRBJ01-01 | 77 | *01:03/*02:01 | *104:01/*11:01 | *02:01/*05:05 | *02:02/*03:01 | *07:01/*13:03 | *01:01 | *01:01 | - |
| | P1 | CMV lysate | CASKGSTEAFF | TRBV19-01 | TRBJ01-01 | 49 | *01:03/*02:01 | *104:01/*11:01 | *02:01/*05:05 | *02:02/*03:01 | *07:01/*13:03 | *01:01 | *01:01 | - |
| | P1 | CMV lysate | CASKGSTEAFF | TRBV19-01 | TRBJ01-01 | 25 | *01:03/*02:01 | *104:01/*11:01 | *02:01/*05:05 | *02:02/*03:01 | *07:01/*13:03 | *01:01 | *01:01 | - |
| | P1 | CMV lysate | CASKGSTEAFF | TRBV19-01 | TRBJ01-01 | 7 | *01:03/*02:01 | *104:01/*11:01 | *02:01/*05:05 | *02:02/*03:01 | *07:01/*13:03 | *01:01 | *01:01 | - |
| | P1 | CMV lysate | CASKGSTEAFF | TRBV19-01 | TRBJ01-01 | 3 | *01:03/*02:01 | *104:01/*11:01 | *02:01/*05:05 | *02:02/*03:01 | *07:01/*13:03 | *01:01 | *01:01 | - |
| | P4 | CMV lysate | CASKGSTEAFF | TRBV19-01 | TRBJ01-01 | 28 | *01:03/*04:01 | - | *01:02/*06:01 | *03:01/*06:02 | *08:03/*15:01 | - | - | *01:01 |
| | P4 | CMV lysate | CASKGSTEAFF | TRBV19-01 | TRBJ01-01 | 1 | *01:03/*04:01 | - | *01:02/*06:01 | *03:01/*06:02 | *08:03/*15:01 | - | - | *01:01 |
| | P9 | CMV lysate | CASKGSTEAFF | TRBV19-01 | TRBJ01-01 | 93 | - | *04:01/*04:01 | *01:01/*06:01 | *03:01/*05:01 | *01:03/*08:03 | - | - | - |
| | P9 | CMV lysate | CASKGSTEAFF | TRBV19-01 | TRBJ01-01 | 12 | - | *04:01/*04:01 | *01:01/*06:01 | *03:01/*05:01 | *01:03/*08:03 | - | - | - |
| | P9 | CMV lysate | CASKGSTEAFF | TRBV19-01 | TRBJ01-01 | 2 | - | *04:01/*04:01 | *01:01/*06:01 | *03:01/*05:01 | *01:03/*08:03 | - | - | - |
| | P9 | CMV lysate | CASKGSTEAFF | TRBV19-01 | TRBJ01-01 | 2 | - | *04:01/*04:01 | *01:01/*06:01 | *03:01/*05:01 | *01:03/*08:03 | - | - | - |
| | P9 | CMV lysate | CASKGSTEAFF | TRBV19-01 | TRBJ01-01 | 2 | - | *04:01/*04:01 | *01:01/*06:01 | *03:01/*05:01 | *01:03/*08:03 | - | - | - |
| | P9 | CMV lysate | CASKGSTEAFF | TRBV19-01 | TRBJ01-01 | 1 | - | *04:01/*04:01 | *01:01/*06:01 | *03:01/*05:01 | *01:03/*08:03 | - | - | - |
| | P1 | CMV lysate | CATAGSTEAFF | TRBV19-01 | TRBJ01-01 | 3 | *01:03/*02:01 | *104:01/*11:01 | *02:01/*05:05 | *02:02/*03:01 | *07:01/*13:03 | *01:01 | *01:01 | - |
| | P9 | CMV lysate | CATAGSTEAFF | TRBV19-01 | TRBJ01-01 | 30 | - | *04:01/*04:01 | *01:01/*06:01 | *03:01/*05:01 | *01:03/*08:03 | - | - | - |
| | P9 | CMV lysate | CAACGSTEAFF | TRBV19-01 | TRBJ01-01 | 1 | - | *04:01/*04:01 | *01:01/*06:01 | *03:01/*05:01 | *01:03/*08:03 | - | - | - |
| Group 2 - CMV- SIGG%GGYE - | P4 | CMV lysate | CASSIGGTGGYEQYF | TRBV19-01 | TRBJ02-07 | 21 | *01:03/*04:01 | - | *01:02/*06:01! | *03:01/*06:02 | *08:03/*15:01 | - | - | *01:01 |
| | P9 | CMV lysate | CASSIGGTGGYEQYF | TRBV19-01 | TRBJ02-07 | 10 | - | *04:01/*04:01 | *01:01/*06:01! | *03:01/*05:01 | *01:03/*08:03 | - | - | - |

| | | | | | | | | | | | | | | |
|---|---|---|---|---|---|---|---|---|---|---|---|---|---|---|
| **global** | P9 | CMV lysate | CASSIGGTGGYEQYF | TRBV19-01 | TRBJ02-07 | 8 | - | *04:01/*04:01 | *01:01/*06:01! | *03:01/*05:01 | *01:03/*08:03 | - | - | - |
| | P9 | CMV lysate | CASSIGGSGGYEQYF | TRBV19-01 | TRBJ02-07 | 84 | - | *04:01/*04:01 | *01:01/*06:01! | *03:01/*05:01 | *01:03/*08:03 | - | - | - |
| | P9 | CMV lysate | CASSIGGSGGYEQYF | TRBV19-01 | TRBJ02-07 | 31 | - | *04:01/*04:01 | *01:01/*06:01! | *03:01/*05:01 | *01:03/*08:03 | - | - | - |
| | P4 | CMV lysate | CASSIGGQGGYEQYF | TRBV19-01 | TRBJ02-07 | 5 | *01:03/*04:01 | - | *01:02/*06:01! | *03:01/*06:02 | *08:03/*15:01 | - | - | *01:01 |
| | P9 | CMV lysate | CASSIGGNGGYEQYF | TRBV19-01 | TRBJ02-07 | 3 | - | *04:01/*04:01 | *01:01/*06:01! | *03:01/*05:01 | *01:03/*08:03 | - | - | - |
| | P4 | CMV lysate | CASSIGGAGGYEQYF | TRBV19-01 | TRBJ02-07 | 1 | *01:03/*04:01 | - | *01:02/*06:01! | *03:01/*06:02 | *08:03/*15:01 | - | - | *01:01 |
| **Group 3 - CMV - SLTGG%SGNT - global** | P2 | CMV lysate | CASSLTGGASGNTIYF | TRBV07-03 | TRBJ01-03 | 5 | *01:03/*02:01 | *02:01/*17:01 | *01:01/*01:02 | *05:01/*06:09 | *01:02/*13:02 | *03:01 | - | - |
| | P2 | CMV lysate | CASSLTGGASGNTIYF | TRBV07-03 | TRBJ01-03 | 2 | *01:03/*02:01 | *02:01/*17:01 | *01:01/*01:02 | *05:01/*06:09 | *01:02/*13:02 | *03:01 | - | - |
| | P2 | CMV lysate | CASSLTGGSSGNTIYF | TRBV07-03 | TRBJ01-03 | 10 | *01:03/*02:01 | *02:01/*17:01 | *01:01/*01:02 | *05:01/*06:09 | *01:02/*13:02 | *03:01 | - | - |
| | P2 | CMV lysate | CASSLTGGSSGNTIYF | TRBV07-03 | TRBJ01-03 | 4 | *01:03/*02:01 | *02:01/*17:01 | *01:01/*01:02 | *05:01/*06:09 | *01:02/*13:02 | *03:01 | - | - |
| | P2 | CMV lysate | CASSLTGGNSGNTIYF | TRBV07-03 | TRBJ01-03 | 106 | *01:03/*02:01 | *02:01/*17:01 | *01:01/*01:02 | *05:01/*06:09 | *01:02/*13:02 | *03:01 | - | - |
| | P2 | CMV lysate | CASSLTGGNSGNTIYF | TRBV07-03 | TRBJ01-03 | 57 | *01:03/*02:01 | *02:01/*17:01 | *01:01/*01:02 | *05:01/*06:09 | *01:02/*13:02 | *03:01 | - | - |
| | P2 | CMV lysate | CASSLTGGNSGNTIYF | TRBV07-03 | TRBJ01-03 | 3 | *01:03/*02:01 | *02:01/*17:01 | *01:01/*01:02 | *05:01/*06:09 | *01:02/*13:02 | *03:01 | - | - |
| | P2 | CMV lysate | CASSLTGGNSGNTIYF | TRBV07-03 | TRBJ01-03 | 3 | *01:03/*02:01 | *02:01/*17:01 | *01:01/*01:02 | *05:01/*06:09 | *01:02/*13:02 | *03:01 | - | - |
| | P3 | CMV lysate | CASSLTGGNSGNTIYF | TRBV07-02 | TRBJ01-03 | 30 | *02:01 | *01:01/*11:01 | *01:01/*01:02 | *05:01/*06:02 | *01:02/*15:01 | - | - | *01:01 |
| | P3 | CMV lysate | CASSLTGGNSGNTIYF | TRBV07-03 | TRBJ01-03 | 16 | *02:01 | *01:01/*11:01 | *01:01/*01:02 | *05:01/*06:02 | *01:02/*15:01 | - | - | *01:01 |
| | P3 | CMV lysate | CASSLTGGNSGNTIYF | TRBV07-02 | TRBJ01-03 | 12 | *02:01 | *01:01/*11:01 | *01:01/*01:02 | *05:01/*06:02 | *01:02/*15:01 | - | - | *01:01 |
| | P3 | CMV lysate | CASSLTGGNSGNTIYF | TRBV07-03 | TRBJ01-03 | 10 | *02:01 | *01:01/*11:01 | *01:01/*01:02 | *05:01/*06:02 | *01:02/*15:01 | - | - | *01:01 |
| | P3 | CMV lysate | CASSLTGGNSGNTIYF | TRBV07-02 | TRBJ01-03 | 5 | *02:01 | *01:01/*11:01 | *01:01/*01:02 | *05:01/*06:02 | *01:02/*15:01 | - | - | *01:01 |
| | P3 | CMV lysate | CASSLTGGNSGNTIYF | TRBV07-02 | TRBJ01-03 | 5 | *02:01 | *01:01/*11:01 | *01:01/*01:02 | *05:01/*06:02 | *01:02/*15:01 | - | - | *01:01 |
| | P3 | CMV lysate | CASSLTGGNSGNTIYF | TRBV07-03 | TRBJ01-03 | 3 | *02:01 | *01:01/*11:01 | *01:01/*01:02 | *05:01/*06:02 | *01:02/*15:01 | - | - | *01:01 |
| | P3 | CMV lysate | CASSLTGGNSGNTIYF | TRBV07-02 | TRBJ01-03 | 3 | *02:01 | *01:01/*11:01 | *01:01/*01:02 | *05:01/*06:02 | *01:02/*15:01 | - | - | *01:01 |
| | P3 | CMV lysate | CASSLTGGNSGNTIYF | TRBV07-03 | TRBJ01-03 | 2 | *02:01 | *01:01/*11:01 | *01:01/*01:02 | *05:01/*06:02 | *01:02/*15:01 | - | - | *01:01 |
| | P3 | CMV lysate | CASSLTGGNSGNTIYF | TRBV07-02 | TRBJ01-03 | 2 | *02:01 | *01:01/*11:01 | *01:01/*01:02 | *05:01/*06:02 | *01:02/*15:01 | - | - | *01:01 |
| | P3 | CMV lysate | CASSLTGGNSGNTIYF | TRBV07-03 | TRBJ01-03 | 2 | *02:01 | *01:01/*11:01 | *01:01/*01:02 | *05:01/*06:02 | *01:02/*15:01 | - | - | *01:01 |
| | P3 | CMV lysate | CASSLTGGNSGNTIYF | TRBV07-03 | TRBJ01-03 | 1 | *02:01 | *01:01/*11:01 | *01:01/*01:02 | *05:01/*06:02 | *01:02/*15:01 | - | - | *01:01 |
| | P3 | CMV lysate | CASSLTGGNSGNTIYF | TRBV07-02 | TRBJ01-03 | 1 | *02:01 | *01:01/*11:01 | *01:01/*01:02 | *05:01/*06:02 | *01:02/*15:01 | - | - | *01:01 |
| | P3 | CMV lysate | CASSLTGGNSGNTIYF | TRBV07-03 | TRBJ01-03 | 1 | *02:01 | *01:01/*11:01 | *01:01/*01:02 | *05:01/*06:02 | *01:02/*15:01 | - | - | *01:01 |
| | P3 | CMV lysate | CASSLTGGNSGNTIYF | TRBV07-03 | TRBJ01-03 | 1 | *02:01 | *01:01/*11:01 | *01:01/*01:02 | *05:01/*06:02 | *01:02/*15:01 | - | - | *01:01 |
| | P2 | CMV lysate | CASSLTGGDSGNTIYF | TRBV07-03 | TRBJ01-03 | 7 | *01:03/*02:01 | *02:01/*17:01 | *01:01/*01:02 | *05:01/*06:09 | *01:02/*13:02 | *03:01 | - | - |
| **Group 4 - Gag - SPP%GE - global** | P3 | Gag pool | CASSPPSGELFF | TRBV18-01 | TRBJ02-02 | 8 | *02:01 | *01:01/*11:01 | *01:01/*01:02 | *05:01/*06:02 | *01:02/*15:01 | - | - | *01:01 |
| | P3 | Gag pool | CASSPPSGELFF | TRBV18-01 | TRBJ02-02 | 7 | *02:01 | *01:01/*11:01 | *01:01/*01:02 | *05:01/*06:02 | *01:02/*15:01 | - | - | *01:01 |
| | P3 | Gag pool | CASSPPSGELFF | TRBV18-01 | TRBJ02-02 | 1 | *02:01 | *01:01/*11:01 | *01:01/*01:02 | *05:01/*06:02 | *01:02/*15:01 | - | - | *01:01 |
| | P3 | Gag pool | CASSPPSGELFF | TRBV18-01 | TRBJ02-02 | 1 | *02:01 | *01:01/*11:01 | *01:01/*01:02 | *05:01/*06:02 | *01:02/*15:01 | - | - | *01:01 |
| | P4 | Gag pool | CASSPPSGELFF | TRBV18-01 | TRBJ02-02 | 9 | *01:03/*04:01 | - | *01:02/*06:01 | *03:01/*06:02 | *08:03/*15:01 | - | - | *01:01 |
| | P2 | Gag pool | CASSPPSGEQYF | TRBV18-01 | TRBJ02-07 | 1 | *01:03/*02:01 | *02:01/*17:01 | *01:01/*01:02 | *05:01/*06:09 | *01:02/*13:02 | *03:01 | - | - |
| | P3 | Gag pool | CASSPPGGEQYV | TRBV18-01 | TRBJ02-07 | 1 | *02:01 | *01:01/*11:01 | *01:01/*01:02 | *05:01/*06:02 | *01:02/*15:01 | - | - | *01:01 |
| | P3 | Gag pool | CASSPPGGEQFF | TRBV07-06 | TRBJ02-01 | 1 | *02:01 | *01:01/*11:01 | *01:01/*01:02 | *05:01/*06:02 | *01:02/*15:01 | - | - | *01:01 |
| | P3 | Gag pool | CASSPPTGELFF | TRBV18-01 | TRBJ02-02 | 17 | *02:01 | *01:01/*11:01 | *01:01/*01:02 | *05:01/*06:02 | *01:02/*15:01 | - | - | *01:01 |
| | P3 | Gag pool | CASSPPTGELFF | TRBV18-01 | TRBJ02-02 | 2 | *02:01 | *01:01/*11:01 | *01:01/*01:02 | *05:01/*06:02 | *01:02/*15:01 | - | - | *01:01 |
| | P3 | Gag pool | CASSPPTGELFF | TRBV18-01 | TRBJ02-02 | 1 | *02:01 | *01:01/*11:01 | *01:01/*01:02 | *05:01/*06:02 | *01:02/*15:01 | - | - | *01:01 |
| | P3 | Gag pool | CASSPPTGELFF | TRBV18-01 | TRBJ02-02 | 1 | *02:01 | *01:01/*11:01 | *01:01/*01:02 | *05:01/*06:02 | *01:02/*15:01 | - | - | *01:01 |
| | P5 | Gag pool | CASSPPTGELFF | TRBV18-01 | TRBJ02-02 | 1 | *02:01 | *01:01 | *02:01/*04:01 | *02:02/*03:19 | *07:01/*08:04 | - | *01:03 | - |
| | P4 | Gag pool | CASSPPGGELFF | TRBV18-01 | TRBJ02-02 | 37 | *01:03/*04:01 | - | *01:02/*06:01 | *03:01/*06:02 | *08:03/*15:01 | - | - | *01:01 |
| | P4 | Gag pool | CASSPPGGELFF | TRBV18-01 | TRBJ02-02 | 30 | *01:03/*04:01 | - | *01:02/*06:01 | *03:01/*06:02 | *08:03/*15:01 | - | - | *01:01 |
| | P4 | Gag pool | CASSPPGGELFF | TRBV18-01 | TRBJ02-02 | 18 | *01:03/*04:01 | - | *01:02/*06:01 | *03:01/*06:02 | *08:03/*15:01 | - | - | *01:01 |
| | P4 | Gag pool | CASSPPGGELFF | TRBV18-01 | TRBJ01-04 | 9 | *01:03/*04:01 | - | *01:02/*06:01 | *03:01/*06:02 | *08:03/*15:01 | - | - | *01:01 |
| | P4 | Gag pool | CASSPPGGELFF | TRBV18-01 | TRBJ02-02 | 1 | *01:03/*04:01 | - | *01:02/*06:01 | *03:01/*06:02 | *08:03/*15:01 | - | - | *01:01 |

**Supplementary Table S5.** Likelihoods that clones would reach the observed sizes under a model of homeostatic proliferation.

| Clone ID | Time since VL <50cpl/mL (years) | Total size of observed clone | Total divisions needed | Proliferation rate (/year) | Net decay (/year) | Half-life (months) | Probability any clone is at least this big | Net decay (/year) | Half-life (months) | Probability any clone is at least this big |
|---|---|---|---|---|---|---|---|---|---|---|
| p1.c.mkl1 | 4.8 | $1.28*10^{7}$ | 23.6 | 10 | -0.19 | 44 | **0** | 0 | ∞ | **0** |
| | | | | 100 | -0.19 | 44 | **0** | 0 | ∞ | **0** |
| | | | | 1000 | -0.19 | 44 | **0** | 0 | ∞ | **0** |
| p1.c.memo1 | 4.8 | $1.42*10^{6}$ | 20.4 | 10 | -0.19 | 44 | **0** | 0 | ∞ | **0** |
| | | | | 100 | -0.19 | 44 | **0** | 0 | ∞ | **0** |
| | | | | 1000 | -0.19 | 44 | **$9*10^{-195}$** | 0 | ∞ | **$2*10^{-127}$** |
| p2.c.stat5b | 11.9 | $1.59*10^{6}$ | 20.6 | 10 | -0.19 | 44 | **0** | 0 | ∞ | **0** |
| | | | | 100 | -0.19 | 44 | **0** | 0 | ∞ | **0** |
| | | | | 1000 | -0.19 | 44 | **$5*10^{-147}$** | 0 | ∞ | **$10^{-57}$** |
| p2.c.hivep3 | 11.9 | $1.18*10^{5}$ | 16.8 | 10 | -0.19 | 44 | **0** | 0 | ∞ | **0** |
| | | | | 100 | -0.19 | 44 | **$18^{-107}$** | 0 | ∞ | **$10^{-41}$** |
| | | | | 1000 | -0.19 | 44 | **$6*10^{-11}$** | 0 | ∞ | **$8*10^{-4}$** |
| p3.c.bach2 | 10.2 | $3.44*10^{5}$ | 18.4 | 10 | -0.19 | 44 | **0** | 0 | ∞ | **0** |
| | | | | 100 | -0.19 | 44 | **0** | 0 | ∞ | **$5*10^{-145}$** |
| | | | | 1000 | -0.19 | 44 | **$4*10^{-33}$** | 0 | ∞ | **$4*10^{-14}$** |
| p3.c.fxbo22 | 10.2 | $8.52*10^{5}$ | 19.7 | 10 | -0.19 | 44 | **0** | 0 | ∞ | **0** |
| | | | | 100 | -0.19 | 44 | **0** | 0 | ∞ | **0** |
| | | | | 1000 | -0.19 | 44 | **$5*10^{-84}$** | 0 | ∞ | **$10^{-35}$** |
| p4.c.st6 | 8 | $2.40*10^{7}$ | 24.5 | 10 | -0.19 | 44 | **0** | 0 | ∞ | **0** |
| | | | | 100 | -0.19 | 44 | **0** | 0 | ∞ | **0** |
| | | | | 1000 | -0.19 | 44 | **0** | 0 | ∞ | **0** |
| p5.c.kcnc2 | 6.5 | $4.95*10^{7}$ | 25.6 | 10 | -0.19 | 44 | **0** | 0 | ∞ | **0** |
| | | | | 100 | -0.19 | 44 | **0** | 0 | ∞ | **0** |
| | | | | 1000 | -0.19 | 44 | **0** | 0 | ∞ | **0** |
| p5.c.delec1 | 6.5 | $3.19*10^{6}$ | 21.6 | 10 | -0.19 | 44 | **0** | 0 | ∞ | **0** |
| | | | | 100 | -0.19 | 44 | **0** | 0 | ∞ | **0** |
| | | | | 1000 | -0.19 | 44 | **0** | 0 | ∞ | **$3*10^{-212}$** |

**Supplementary Table S7.** High resolution HLA genotyping of the study participants.

High resolution HLA typing of participants included in the TCR sequence analyses

| Participant ID | A | C | B | DRB1 | DRB3 | DRB4 | DRB5 | DQA1 | DQB1 | DPA1 | DPB1 |
|---|---|---|---|---|---|---|---|---|---|---|---|
| P1 | *02:01 | *17:03 | *41:02 | *07:01 | *01:01 | *01:01 | | *02:01 | *02:02 | *01:03 | *11:01 |
| | *29:02 | *16:01 | *44:03 | *13:03 | | | | *05:05 | *03:01 | *02:01 | *104:01 |
| P2 | *02:01 | *08:02 | *14:02 | *01:02 | *03:01 | | | *01:01 | *05:01 | *01:03 | *02:01 |
| | *03:01 | *07:01 | *49:01 | *13:02 | | | | *01:02 | *06:09 | *02:01 | *17:01 |
| P3 | *02:01 | *01:02 | *27:05 | *01:02 | | | *01:01 | *01:01 | *05:01 | *02:01 | *01:01 |
| | *68:01 | *04:01 | *53:01 | *15:01 | | | | *01:02 | *06:02 | | *11:01 |
| P4 | *02:01 | *07:02 | *07:02 | *08:03 | | | *01:01 | *06:01 | *03:01 | *01:03 | *104:01 |
| | *68:01 | *03:04 | *40:01 | *15:01 | | | | *01:02 | *06:02 | | |
| P5 | *30:01 | *04:01 | *53:01 | *07:01 | | *01:03 | | *02:01 | *02:02 | *02:01 | *01:01 |
| | *30:02 | *07:01 | *57:03 | *08:04 | | | | *04:01 | *03:19 | | |
| P6 | *26:01 | *04:01 | *35:01 | *11:01 | *02:02 | | | *01:02 | *06:02 | *02:01 | *01:01 |
| | *30:02 | *18:01 | *81:01 | *13:04 | | | | *05:05 | *03:19 | | *11:01 |
| P8 | *03:01 | *07:02 | *07:02 | *07:01 | | *01:01 | *01:01 | *02:01 | *02:02 | *01:03 | *02:02 |
| | | *07:01 | *08:01 | *15:01 | | | | *01:02 | *06:02 | | *03:01 |
| P9 | *11:01 | *04:01 | *35:01 | *08:03 | | | | *01:01 | *05:01 | | *04:01 |
| | | *15:02 | *51:01 | *01:03 | | | | *06:01 | *03:01 | | |
